# Decoding Agency-Related Neural States During Human–AI Interaction in Autonomous Driving Using EEG and Deep Learning

**DOI:** 10.64898/2026.07.22.740014

**Authors:** Eléonore Houdoyer, Solène Le Bars, Valérian Chambon

## Abstract

The sense of agency, the experience of controlling one’s actions and their consequences, is a fundamental component of human interaction with autonomous systems. As artificial intelligence increasingly mediates decision-making in domains such as autonomous driving, understanding and monitoring agency-related processes becomes critical for maintaining user engagement, trust, and appropriate levels of control. However, existing approaches to measuring agency rely primarily on subjective reports or event-based neural markers, which are poorly suited for naturalistic and continuous human–AI interaction. In this study, we investigate whether agency-related neural states can be decoded from ongoing electroencephalographic (EEG) activity using deep learning. Across two experiments, we manipulated agency through (i) decision authority (human vs AI control) and (ii) system explainability (AI with vs without intention-based explanations). Behavioral results confirmed that both manipulations significantly modulated participants’ perceived control. At the neural level, we trained EEGNet-based models to decode agency-modulating experimental conditions from pre-feedback EEG activity. Decoding performance was robust at the intra-subject level and remained significantly above chance across participants using a leave-one-subject-out framework, demonstrating partial cross-subject generalization of agency-related neural representations. Spectral ablation analyses revealed that low-frequency activity, particularly in the delta and theta bands, made the dominant contribution to decoding performance. Complementary time–frequency analyses showed that these bands exhibited increased power under reduced-agency conditions, specifically during the post-keypress, pre-feedback interval. Together, these findings indicate that agency-related information is embedded in continuous, low-frequency neural dynamics associated with predictive monitoring processes. By demonstrating that such information can be decoded from single-trial EEG in an offline setting, this work provides a foundation for future real-time, non-intrusive monitoring of user states in human–AI interaction. These results open new avenues for the development of neuroadaptive systems capable of dynamically regulating automation and explainability to preserve human agency.

## 1. Introduction

Artificial intelligence (AI) systems are increasingly embedded in everyday environments, mediating a wide range of human activities such as driving, decision support, and adaptive control. In many of these contexts, humans no longer exert direct control over system behavior but instead interact with autonomous agents that perceive, decide, and act on their behalf. As a result, interaction with AI systems has shifted from direct manipulation toward supervisory and collaborative forms of control (Pagliari et al.). This shift has positioned human–AI interaction (HAI) at the intersection of cognitive neuroscience and artificial intelligence, raising fundamental questions about how humans experience control, responsibility, and engagement when interacting with intelligent systems.

A central construct for understanding human experience in automated and human-AI interaction is the sense of agency (SoA), defined as the subjective experience of being the author of one’s actions and their consequences (Chambon & Haggard., 2012). The sense of agency supports goal-directed behavior by enabling responsibility attribution, sustained engagement, and effective monitoring of external systems. Converging evidence indicates that increasing levels of automation tend to attenuate this experience (Endsley & Kiris, 1995; Kaber & Endsley, 1997), leading to disengagement and so-called out-of-the-loop behavior, particularly in safety-critical domains (Ephrath & Young, 1981; Kessel & Wickens, 1982). Understanding how agency is altered, and potentially restored, during interaction with AI systems therefore constitutes a key challenge for both cognitive science and human-centered AI design. From a cognitive perspective, agency is commonly conceptualized as emerging from predictive processes that operate across multiple levels of intentional representation (Pacherie,2008).

Human intentions are hierarchically structured, encompassing action-related plans, intermediate subgoals at a proximal level, and more abstract, long-term goals at a distal level (Chambon et al., 2011). This hierarchical organization supports the anticipation of action outcomes and enables the continuous integration of sensory feedback during action control (Pacherie, 2008, 2014). In human joint action, the ability to represent and anticipate a partner’s intentions at both proximal and distal levels allows interacting agents to coordinate their behavior and align predictions over time, thereby preserving their sense of agency (van der Wel 2015; van der Wel et al. 2012; Le Bars et al.,2020,2022).

By contrast, during interaction with artificial agents, this form of intentional alignment and prediction is often disrupted by the opacity of algorithmic decision-making. When users are unable to infer an agent’s goals or action plans, the predictive mechanisms that support control and monitoring are compromised, resulting in diminished agency and reduced engagement (Sahaï et al., 2019; Berberian, 2019; Vantrepotte et al., 2022). This observation has motivated the development of explainable artificial intelligence (XAI), which aims to restore transparency and predictability by providing users with information about artificial agents’ goals, decisions, or action-level processes.

Building on principles from human joint action, intention-based explainability has been proposed as a particularly promising approach to preserving the sense of agency during human-AI interaction. Behavioral and neurophysiological studies, including ERP-based investigations, have shown that increased AI intervention tends to reduce the sense of agency (Berberian, 2019; Vantrepotte et al., 2022; Houdoyer et al., under review), whereas enabling artificial agents to disclose and explain their intentions can restore it (Le Goff et al.,2018; Houdoyer et al., under review). Together, these findings provide converging evidence that intention sharing constitutes an effective mechanism for supporting agency in human-AI interaction.

However, despite these established effects, existing studies have primarily relied on experimental paradigms in which agency is assessed through temporally isolated action–outcome events (Le Goff et al., 2018; Vantrepotte et al., 2022; Houdoyer et al., under review). This leaves open the question of how agency-related processes unfold and can be monitored during extended phases of continuous interaction, such as ongoing system-controlled behavior. Accordingly, the critical challenge is no longer whether intention-based explainability can modulate the sense of agency, but whether such effects can be tracked under interaction conditions that more closely reflect everyday human-AI interaction, in which control is continuous, system behavior evolves dynamically, and clear action–outcome boundaries are often absent.

This challenge is closely tied to how the sense of agency is currently measured. Explicit measures, such as self-reported ratings of control or responsibility, are straightforward to administer but are highly susceptible to subjective biases and demand characteristics (Tsakiris & Haggard, 2005; Wen et al.,2018). Moreover, their repeated administration disrupts ongoing interaction, limiting ecological validity and precluding their use in continuous or adaptive human–AI systems (Wen et al.,2018). Implicit behavioral measures, including intentional binding and sensory attenuation, provide more objective indices of agency but fundamentally rely on discrete, well-isolated action–outcome events and require extensive averaging across trials. Consequently, they are ill-suited for capturing agency as a continuously evolving process (Wen et al.,2018).

Neurophysiological approaches face similar limitations. Event-related potential (ERP) studies have identified reliable neural correlates of agency, such as attenuations of early sensory components (e.g., N1 attenuation) and modulations of later stages of outcome evaluation (e.g., increases in the P2 component) under conditions associated with higher levels of agency (Houdoyer et al., in prep.; Kühn et al., 2011;Timm et al., 2014; Bednark et al., 2015). While these findings establish a clear link between agency and measurable neural dynamics, ERP-based methods inherently depend on event locking and trial averaging, which limits their applicability in naturalistic interaction contexts characterized by continuous action–feedback loops and overlapping neural responses.

Taken together, these constraints highlight the need for neural measurement frameworks capable of capturing agency-related information continuously and without reliance on predefined events. In this context, time-frequency representations of EEG signals provide access to ongoing, non-phase-locked neural dynamics that unfold over extended timescales and may better reflect the continuous nature of the sense of agency during human-AI interaction.

Within this framework, oscillatory activity in sensorimotor and fronto-parietal networks offers several candidate neural mechanisms relevant to agency-related processes. Alpha–mu rhythms (approximately 8–13 Hz), particularly over central regions, have been associated with sensorimotor gating and the regulation of cortical excitability (Kang et al., 2015; Wen et al., 2017) and typically exhibit sustained desynchronization during voluntary action and action monitoring processes closely linked to perceived control (Kühn et al., 2011; Haggard, 2017). Beta-band activity (approximately 13–30 Hz), often implicated in the maintenance and updating of sensorimotor predictions, shows characteristic modulations during movement execution and feedback processing, making it a plausible marker of predictive control mechanisms underlying agency. In addition, theta-band oscillations (approximately 4–7 Hz), especially over frontal and midline regions, have been linked to performance monitoring (Cavanagh & Frank, 2014), error detection, and the processing of mismatches between expected and actual outcomes (Arnal & Giraud, 2012), processes that are central to the experience of agency (Haggard, 2017).

However, evidence for robust oscillatory markers uniquely attributable to variations in agency during continuous interaction remains limited, suggesting that agency-related information may be distributed across frequency bands, spatial locations, and time, rather than being captured by isolated spectral features.

Recent advances in machine learning, and deep learning in particular, offer new opportunities to address these challenges. Convolutional neural networks (CNNs) have demonstrated a strong capacity to learn hierarchical spatiotemporal representations directly from high-dimensional and noisy neurophysiological data, including raw or minimally preprocessed EEG (Lawhern et al., 2018). By integrating information across channels, time, and spectral content in an end-to-end manner, CNN-based approaches enable single-trial decoding and are well-suited for tracking latent cognitive states that evolve continuously over time (Lawhern et al., 2018).

Building on this perspective, the present work investigates whether neural signatures associated with experimentally induced variations in agency-related processing can be discriminated from ongoing EEG activity during human-AI interaction. Using EEGNet, a compact CNN architecture tailored to electrophysiological data, we implemented a two-stage decoding strategy. First, we assessed whether neural decoding models could distinguish between non-automated (human-controlled) and automated (AI-controlled) interaction while task demands and parameters were highly similar (i.e., number of key presses, visual stimuli etc.). Second, focusing exclusively on automated interaction, we examined whether decoding models could discriminate between AI systems operating with and without intention-based explanatory information.

Beyond decoding performance, we investigated the spectral dynamics underlying successful classification by combining frequency-band ablation analyses with time-frequency decomposition of EEG activity. These complementary approaches allowed us to identify the oscillatory components that contribute most strongly to the discrimination between experimental conditions, thereby highlighting candidate neural mechanisms associated with variations in perceived agency.

Finally, we evaluated decoding performance both within individuals (intra-subject) and across individuals (inter-subject). Intra-subject decoding evaluates whether agency-related neural states can be reliably identified at the individual level, supporting personalized applications at the cost of subject-specific calibration. Inter-subject decoding tests whether learned neural representations generalize across users, a key requirement for scalable human-AI systems that must operate without individual retraining. This distinction directly links neural decoding performance to the practical constraints of deploying agency-aware systems in real-world settings.

By integrating EEG-based neural decoding with a theoretically grounded account of the sense of agency, this work contributes to both flied of cognitive neuroscience and applied artificial intelligence. From a neuroscientific perspective, it provides a computationally informed framework for studying agency-related mechanisms as dynamic and continuously evolving neural processes that extend beyond traditional event-based analyses. From an HAI perspective, it advances the development of agency-aware systems by demonstrating how users’ internal states can be monitored non-intrusively through learned neural representations, thereby enabling adaptive automation and explainability strategies that preserve meaningful human involvement in continuous human-AI interaction.

## 2. Experimental framework

### 2.1 Participants

Two independent experiments were conducted on separate participant cohorts, yielding a total sample of 58 adult volunteers before data quality control. All participants reported normal hearing and normal or corrected-to-normal vision, and declared no history of neurological or psychiatric disorders. Participants were naïve to the aims of the study and provided written informed consent prior to participation. Participant inclusion was determined exclusively based on EEG signal quality, following inspection of the recorded data.

Datasets exhibiting substantial noise, predominantly prolonged low-frequency drifts, were excluded when such artefacts resulted in an insufficient number of analysable trials. Specifically, participants contributing fewer than 40 valid trials in the experimental conditions of interest were excluded from further analyses. This quality-control procedure was defined a priori and applied prior to any decoding or condition-specific analyses, independently of behavioural or neural outcome measures.

After quality control, the final analysed sample comprised 20 participants in Experiment 1 (10 females; mean age = 23.57 ± 3.34 years) and 17 participants in Experiment 2 (8 females; mean age = 23.07 ± 3.26 years). Participants received a fixed compensation of €15. To maintain engagement throughout the task, they were informed that an additional performance-based bonus would be awarded if their score ranked among the top three participants. The amount of this bonus was not specified in advance. In practice, all participants received a €5 bonus at the end of the experiment, resulting in a total compensation of €20. Participants were debriefed about the bonus procedure after completing the study.

All experimental procedures were approved by the local ethics committee (IRB00003888) and were conducted in accordance with the principles of the Declaration of Helsinki.

### 2.2 Apparatus and Materials

All experiments were conducted in a sound-attenuated chamber (Ecole normale supérieure-PSL), designed to minimize ambient acoustic noise and electromagnetic interference. Task presentation and experimental control were implemented using PsychoPy (version 2025.1.1), with event markers synchronized via the Lab Streaming Layer (PyLSL version 1.14) to ensure precise temporal alignment between stimulus presentation, behavioral responses, and EEG acquisition.

Visual stimuli were presented in full-screen mode on a 14-inch LCD monitor (1920 × 1080 pixels; 60 Hz refresh rate), viewed from an approximate distance of 60 cm against a uniform gray background. Auditory feedback was delivered through closed-back headphones (Sennheiser HD 280 Pro), calibrated to an intensity of 80 dB SPL.

Behavioral responses were collected using a standard computer keyboard. All stimulus delivery, response logging, and event synchronization were controlled via custom Python scripts to ensure reproducibility and precise temporal control across experimental sessions

### 2.3 Task Overview and Trial Structure

Participants completed a continuous navigation task, adapted from Houdoyer et al. (under review), in which they either acted directly on the environment or interacted with a belief-based automated assistance system. The paradigm was designed to vary the extent to which control over strategy selection was retained by the participant or delegated to the system during goal-directed behaviour.

At the beginning of each trial, two visual options were displayed, each corresponding to a distinct navigational strategy. A red heart indicated the safe strategy, whereas a banknote symbol indicated the risky strategy. These strategies differed in their implied risk–reward structure: the safe strategy was associated with lower risk and lower reward, whereas the risky strategy was associated with higher risk and higher reward.

In the Motor/no-AI condition, these contingencies were implemented directly: the safe strategy was associated with a 20% obstacle probability and a 2.5-point reward, whereas the risky strategy was associated with a 50% obstacle probability and a 4-point reward. This manual condition allowed participants to learn and experience the characteristic consequences of each strategy, including differences in speed, risk, and reward, and was intended to establish prior expectations about the outcomes associated with each option.

In the belief-based AI conditions, the apparent risk–reward structure was preserved for participants, but obstacle probability was experimentally controlled. Specifically, the difference in obstacle probability between the two strategies was removed: both strategies were associated with a fixed 10% obstacle rate, while reward magnitude continued to depend on the selected strategy. This manipulation was introduced to standardise the number of usable trials across conditions and to maintain trust in the purported automated assistance system. Obstacle trials were nevertheless retained in the task to keep participants attentive and to preserve the credibility of the risk–reward setting. These trials were not considered trials of theoretical interest and were excluded from the main analyses.

The source of control over strategy selection varied across conditions. In the Motor/no-AI condition, participants chose one of the two options and initiated the trial by pressing the left or right arrow key according to the spatial position of the selected target. In the belief-based AI conditions, participants were told that the system selected the strategy autonomously; their role was to initiate the trial by pressing the space bar after the system’s choice had been presented. Thus, despite differences in control delegation, each trial required a single keypress at onset. This matched the basic motor requirement across conditions and reduced the likelihood that differences in behavioural or neural responses reflected differences in motor execution rather than agency-related processes.

Vehicle movement started immediately after the keypress and lasted 300 ms. Once the vehicle reached the target, the outcome was signalled by an auditory tone. Feedback consisted of 100-ms pure tones delivered at 80 dB SPL, with one tone at 500 Hz and the other at 1000 Hz. On most trials, the tone was consistent with the outcome predicted by the selected strategy. On the remaining trials, the tone violated this prediction, providing an unexpected auditory outcome. The visual display was kept unchanged during the auditory-feedback period and until 600 ms after tone onset, in order to avoid visual transients overlapping with auditory-evoked ERP responses. Each trial then ended with a 700-ms blank interval before the next trial began.

To maintain attention, a subset of trials was presented as catch trials. On these trials, auditory-feedback amplitude was reduced by 50%, and participants were asked to respond as quickly as possible using the same key used to initiate the trial. Brief performance feedback was provided at the block level to encourage sustained engagement.

Sense-of-agency ratings were collected on 20% of trials. On these trials, 1200 ms after tone offset, participants answered the question: “How much did you feel in control over the trial?” using a Likert scale ranging from 1 (no control) to 9 (complete control). These ratings were included to monitor task engagement and to verify that the experimental manipulations produced the expected behavioural effects of automation and explanatory support, as observed in previous work using this paradigm (Houdoyer et al., under review). Because the primary focus of the experiment was electrophysiological, rating trials were kept sparse to avoid disrupting task flow and to preserve EEG data quality. The ratings were therefore used as a behavioural manipulation check rather than as measures intended for trial-by-trial correlation with EEG responses.

### 2.4 Experimental Manipulations

Across experiments, the overall task structure, visual displays, and auditory-feedback contingencies were held constant. Experimental conditions differed only in the level of control delegation and, during automated decision-making, in the availability of AI-provided explanatory information. Trial timing was identical across experiments, except in Experiment 2, where a fixed 600-ms cue period was presented before participants were allowed to initiate vehicle movement.

#### Belief-Based AI Assistance Context

In the belief-based AI conditions, participants were led to believe that an AI-based driving assistance system determined the vehicle’s target and movement. The automated task was framed as a supervisory driving situation, in which participants did not continuously steer the vehicle. Instead, they configured the assistance system before the task, observed its decisions during each trial, and confirmed the outcome once the vehicle had reached its destination.

Before starting the automated blocks, participants were told that the assistance system could select the most appropriate strategy and trajectory on a trial-by-trial basis. The system was introduced as a decision-tree algorithm that evaluated three task-relevant criteria: safety, reward, and driving time. Participants were informed that these criteria could conflict with one another and that the system would arbitrate between them when selecting the vehicle’s behaviour.

To make the interaction more credible, participants first set a risk-tolerance parameter for the purported AI system. This setting corresponded to the obstacle probability they were willing to accept and could vary between 15% and 50%. They were told that conservative settings would favour safer choices at the cost of longer travel times and smaller rewards, whereas higher-risk settings would favour faster and more rewarding choices but increase the likelihood of obstacles. Participants were also informed that their risk setting would be combined with the current trial parameters when the system selected the strategy and trajectory.

For experimental control, however, the vehicle’s behaviour was not generated online by an adaptive algorithm. Strategy and trajectory choices were determined in advance according to predefined, counterbalanced sequences, ensuring that participants were exposed to comparable conditions. Consequently, the risk preference entered by participants did not alter the system’s behaviour during the task. Its purpose was to strengthen the belief that the automated decisions reflected both user-defined preferences and task constraints.

The task should therefore be interpreted as a controlled belief-based simulation of AI-assisted conditional automation. Although the scenario was conceptually inspired by SAE Level 3 automated driving, it did not implement a functional SAE Level 3 system. Rather, it created an experimentally standardised context in which participants believed that part of the driving-related decision process had been delegated to an AI-based assistance system.

#### Experiment 1: Manipulation of the Level of Control Delegation

Experiment 1 examined the effect of control delegation by manipulating whether navigational strategy selection was retained by the participant or delegated to the belief-based automated system. In the participant-controlled condition, participants selected the navigational strategy themselves and initiated the trial by pressing the arrow key corresponding to the spatial position of the chosen option.

In the automated-control condition, strategy selection was presented as being performed by the system, and participants initiated the trial by pressing the space bar (see Figures 1A and 1B). In both conditions, a single explicit motor response was required to initiate the trial. This design equated motor execution demands across conditions, thereby allowing neural differences to be attributed more specifically to the level of control delegation rather than to motor-related activity.

**Figure 1.**
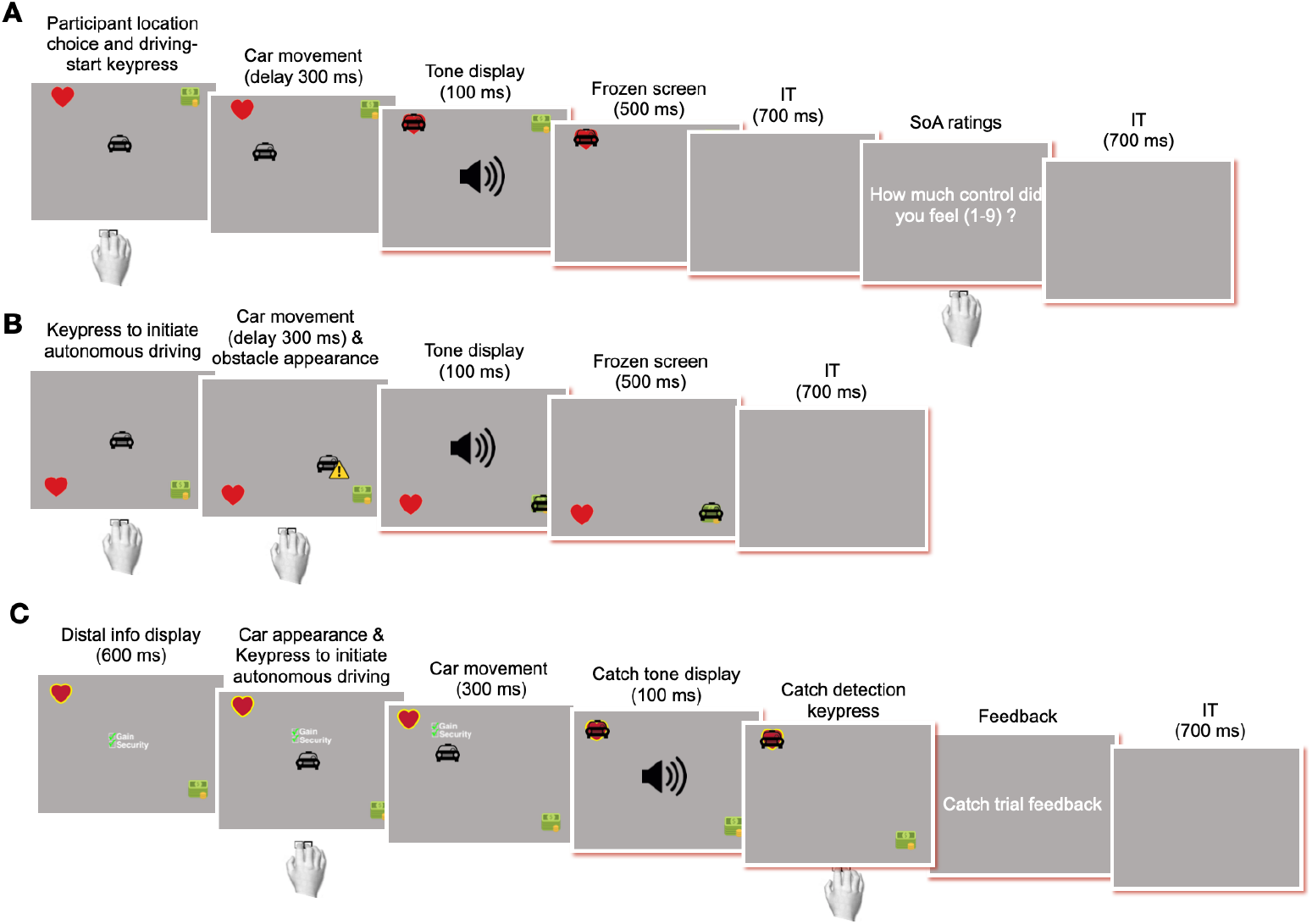
Task layout and trial timeline across motor, AI no-explanation, and AI explanation conditions. Each trial began with the presentation of two targets defining the available strategy options and their associated trajectories. In the motor condition (A), participants selected the target using the left or right arrow key. In the AI conditions (B–C), participants initiated the trial by pressing the space bar, while the AI selected the strategy and trajectory. The car then moved toward the chosen target for 300 ms. In a subset of trials, an obstacle (yellow warning panel) appeared during movement and required a rapid response. Upon target arrival, an auditory tone was presented (match: ∼80%; mismatch/oddball: ∼20%), and on 20% of trials participants rated their sense of agency on a 9-point scale. In Experiment 2, AI trials included a 600-ms pre-movement explanation period. In the no-explanation condition (B), green checkmarks were presented together with semantically uninformative placeholder text. In the AI distal explanation condition (C), the AI-selected strategy was indicated by highlighting the corresponding target and accompanied by green checkmarks and explanatory text. Red-highlighted portions of the trial timeline were excluded from all EEG analyses, including deep-learning decoding and time–frequency analyses.

#### Experiment 2: Explanatory information under AI decision-making

Experiment 2 focused exclusively on belief-based automated decision-making and examined whether providing distal explanatory information modulated neural processing when participants did not control the decision outcome (see Figure 1.B, C).. Participants completed two within-subject conditions, belief-based AI without explanation and belief-based AI with distal explanation, presented in alternating blocks. In both conditions, the purported AI-based assistance system was presented as selecting the strategy by taking into account the task-relevant parameters previously described to participants, namely safety, reward, and driving time, as well as the risk preference configured before the automated block.

Explanations were designed as brief, immediately interpretable cues indicating why the AI-selected strategy was advantageous with respect to these parameters. In the belief-based AI without explanation condition, the display contained green checkmarks and visually matched placeholder text (“Null/Unknown”) presented for 600 ms. This condition provided a control for the presence, timing, and general visual format of explanatory material, while conveying no meaningful information about the AI’s decision. In the belief-based AI with distal explanation condition, the strategy selected by the system was indicated by highlighting the corresponding target. This visual cue provided information about the AI’s goal-level decision, namely the selected strategy, without displaying an additional strategy icon. The highlight was accompanied by brief feature-related statements, such as “Gain” and “Security”, presented with green checkmarks for 600 ms. Thus, the explanation combined a direct visual indication of the AI’s upcoming distal choice with concise textual information about the parameters presented as guiding that choice.

The displays were matched across conditions in timing and overall visual structure so that differences between conditions could be attributed to the informational value of the explanation rather than to the mere presence of additional visual elements. In both conditions, the vehicle appeared only after the 600-ms cue period, indicating that participants were allowed to initiate movement by pressing the space bar. Participants had no influence over the selected strategy, and motor requirements were identical across conditions.

Importantly, the explanatory cues were prospective rather than retrospective: they were presented before movement onset and were intended to support anticipation and monitoring of the AI’s upcoming decision, rather than to provide a post hoc justification of a completed action.

### 2.5 Procedure and Trial Counts

The two experiments were conducted on independent participant cohorts, with no overlap between samples. All preprocessing, decoding, and statistical analyses were performed separately for each experiment.

Within each experiment, participants completed eight blocks of 50 trials per condition, yielding 400 trials per condition. Each block comprised 40 match trials, in which auditory feedback corresponded to the expected outcome, and 10 mismatch trials, in which auditory feedback violated the expected outcome. Mismatch trials therefore represented 20% of trials in each block.

Conditions were presented in alternating blocks throughout the session. In Experiment 1, for example, blocks alternated between the Motor/no-AI and belief-based AI conditions, with the starting condition counterbalanced across participants. In Experiment 2, blocks alternated between the belief-based AI without explanation and belief-based AI with distal explanation conditions, again with the starting condition counterbalanced across participants. This ensured that condition order was balanced across the sample and that potential order effects were controlled.

Obstacle trials were interspersed pseudo-randomly throughout the session. In the belief-based AI conditions, these trials occurred on 10% of trials. In the Motor/no-AI condition, obstacle occurrence depended on the strategy selected by the participant and therefore followed the obstacle probability associated with the chosen strategy rather than a fixed predefined rate ( see 2.3 Task Overview and Trial Structure).. Obstacle trials also served as catch trials: auditory-feedback amplitude was reduced by 50%, and participants were instructed to respond as quickly as possible using the same key used for trial initiation. Catch and obstacle trials were excluded from all decoding analyses.

In addition, 20% of all trials included a sense-of-agency rating, presented 1200 ms after auditory feedback. Participants rated their feeling of control over the trial using the scale described above. These ratings served as a behavioural manipulation check and were not used for trial-level EEG decoding.

The assignment of high- and low-pitched tones to expected and unexpected outcomes was counterbalanced across participants to prevent frequency-specific confounds. Before the main task, participants completed a practice phase consisting of 10 trials per condition to familiarise themselves with the task structure, response mappings, and feedback contingencies. Short breaks were provided between blocks to reduce fatigue and maintain EEG signal quality. After preprocessing and trial rejection, the number of retained trials per participant was sufficient to support within-subject and inter-subject decoding analyses.

### 2.6 EEG Acquisition

Electroencephalographic (EEG) activity was recorded using a Neuroelectrics Enobio 20 system equipped with a custom 17-electrode scalp montage (Fp1, Fpz, Fp2, F7, F3, Fz, F4, F8, C3, Cz, C4, T7, P7, Pz, P8, T8, Oz). Electrode locations were digitized in three-dimensional Cartesian space for each participant to ensure reproducible sensor placement and accurate spatial correspondence across recordings.

Signals were acquired at a sampling rate of 500 Hz using NIC software (version 2.1.3.11). During acquisition, an online 50 Hz notch filter was applied to attenuate power-line interference. No additional online filtering, re-referencing, or artefact correction procedures were applied, such that all signal conditioning decisions were implemented uniformly during offline preprocessing.

EEG amplitudes were recorded in nanovolts. In addition to EEG, a three-axis accelerometer signal was recorded concurrently at 100 Hz to monitor gross head movements; these data were not included in subsequent analyses.

### 2.7 EEG Data Preprocessing

Offline preprocessing was performed using MNE-Python (version 1.11.0) within a predefined, semi-automated processing pipeline applied identically across participants and experiments. All preprocessing steps were specified a priori and executed independently of experimental conditions and decoding analyses.

Raw continuous EEG recordings were first visually inspected to identify and exclude segments containing prominent artefacts. Channels exhibiting abnormal signal characteristics were then identified using a composite approach combining statistical deviation measures and inter-channel correlation metrics, supplemented by automated diagnostics provided by the PyPREP toolbox. Channels identified as unreliable were marked as bad and subsequently reconstructed using spherical spline interpolation.

Following channel cleaning, the data were filtered offline using 50 and 100 Hz notch filters to attenuate residual mains contamination and its first harmonic not fully suppressed by the online 50 Hz notch applied during acquisition. The signal was then band-pass filtered between 1 and 40 Hz using a zero-phase finite impulse response (FIR) filter implemented in MNE-Python. This frequency range was selected as a methodological compromise to retain EEG activity relevant to cognitive and predictive processes while minimizing slow drifts and high-frequency muscle artifacts, thereby optimizing signal quality for deep learning-based single-trial decoding. Although a 1 Hz high-pass filter attenuates the slowest fluctuations, this cutoff was intended to reduce low-frequency non-neural drifts that might otherwise dominate the decoding models. Trials exceeding ±200 µV for experiment 1 and ±150µV for experiment 2 (threshold selection based on grid search, see Supplementary Results, Tables 2 and 5) were automatically rejected.

Independent component analysis (ICA) was not applied, as convolutional neural networks are known to tolerate moderate residual artifacts and can learn to down-weight non-informative components during training. Moreover, avoiding ICA reduced the risk of introducing additional assumptions or distortions into the data prior to decoding.

Preprocessed EEG data were segmented into epochs time-locked to action initiation (keypress). Epochs were restricted to the interval preceding auditory feedback onset to ensure that analyses targeted neural activity related to action preparation, initiation, and control rather than outcome-related sensory processing.Because task timing differed between experiments, epoch duration was adapted accordingly.

In Experiment 1, no enforced pre-keypress delay was present, such that epoch duration was constrained by participants’ reaction times; a relatively short window (−280 to +280 ms) was therefore selected to avoid including unrelated pre-trial activity. In Experiment 2, a fixed 600 ms cue period preceded action initiation while AI-provided information was displayed, permitting the use of a longer epoch extending from −560 to +280 ms relative to keypress. Pre-keypress window lengths were determined using a preliminary grid search conducted on the training data, with window sizes selected to maximize mean validation accuracy across participants. This procedure was defined a priori and implemented independently of all final test-set evaluations (see Supplementary Results section, Tables 1 and 6).

**Table 1.** Intra-subject decoding performance in Experiment 1 (N=20). Accuracy, Precision, Recall, F1-score, and AUC were computed for each participant on strictly held-out test data and averaged across five repeated block-wise train/validation/test partitions to obtain participant-level estimates. Group-level means, standard deviations, and 95% t-based confidence intervals were then computed across participants. Chance level for binary classification was 50%.

| Metric (%) | Mean | SD | 95% CI |
| --- | --- | --- | --- |
| Accuracy | 77.6 | 9.17 | [73.3, 81.9] |
| Precision | 78.8 | 9.05 | [74.60, 83.08] |
| Recall | 78.9 | 8.92 | [74.69, 83.04] |
| F1-score | 77.42 | 9.36 | [73.04, 81.8] |
| AUC | 84.25 | 9.73 | [79.70, 88.8] |
| Chance level | 50 | — | — |

Only regular, successfully completed trials were retained for decoding analyses. Regular trials were defined as trials without obstacle events and without catch-tone detection demands. Trials were considered successfully completed when participants produced a single appropriate initiation response, with no additional keypresses suggesting overshooting or an inappropriate response strategy. Catch trials, obstacle trials, and mismatch trials were excluded to avoid introducing attentional, motor, or outcome-related sensory prediction-error signals that could trivially distinguish between conditions. This restriction ensured that decoding models primarily captured neural activity related to anticipatory and control-related processes, rather than post hoc error detection. No behavioural measures were analysed in the present study.

## 3. Neural decoding framework

### 3.1 Neural network architecture (EEGNet)

The convolutional neural network (CNN) used for EEG-based decoding was derived from the classical EEGNet architecture. The model was designed to capture temporally and spatially structured information in low-channel-count EEG recordings while maintaining a limited number of trainable parameters, thereby reducing overfitting risk in low-data regimes.

The network comprises a sequence of processing stages, including an input layer, a temporal convolution layer, a depthwise spatial convolution layer, two average pooling layers with dropout regularization, a separable convolution layer, a fully connected classification layer, and a softmax output layer.

#### Input layer

The network operates on epoched EEG signals represented as three-dimensional tensors of shape (channels x samples x 1), where channels correspond to the number of EEG electrodes (17), samples denote the number of temporal samples per epoch, and the singleton dimension represents a single input feature map.

#### Temporal convolution (Conv1)

The first convolutional layer performs temporal filtering independently for each EEG channel. It applies *F*_1_ = 16 convolutional filters with kernel size (1 x 125), stride 1, and same padding, without bias terms. The temporal kernel length was set to 125 samples (250 ms at 500 Hz), corresponding to a temporal window equivalent to ∼4 Hz spectral sensitivity. This duration allows the kernels to capture physiologically relevant EEG rhythms while preserving local temporal structure within each epoch. The selected value is consistent with kernel durations commonly used in EEGNet-based architectures after scaling for sampling rate.By restricting convolution to the temporal dimension, this layer learns time- and frequency-sensitive representations while preserving spatial independence across EEG channels. The resulting feature maps are batch-normalized to improve numerical stability and facilitate optimization.

#### Depthwise spatial convolution (Conv2)

To model spatial dependencies across EEG channels, a depthwise convolution is applied to the temporally filtered signals. This layer uses kernels of size (channels x 1) with depth multiplier D = 2, allowing multiple spatial filters to be learned for each temporal feature map. Bias terms are omitted, and a max-norm constraint of 1 is applied to the convolutional weights to limit parameter growth and promote stable learning. The output is batch-normalized and passed through an exponential linear unit (ELU) activation function.

#### Pooling and dropout (Pool1)

An average pooling operation with pool size (1 x 4) reduces temporal resolution and computational complexity while preserving distributed temporal information. At a sampling rate of 500 Hz, this corresponds to a local aggregation window of 8 ms, which maintains physiologically relevant EEG dynamics while performing mild temporal compression. Dropout regularization (rate = 0.3 for intra-subject decoding and 0.15 for intersubject decoding, as the training set is much larger) is subsequently applied.

#### Separable convolution (Conv3)

A separable convolutional layer is then applied to further summarize temporal information with reduced parameter count. This layer employs *F*_2_ = 32 filters with kernel size (1 x 16), stride 1, and same padding, without bias terms. The separable convolution factorizes temporal filtering and channel mixing, improving computational efficiency.

The output is batch-normalized, transformed using an ELU activation function, and downsampled using a second average pooling layer (1 x 4) followed by dropout (rate = 0.3 for intra-subject decoding and 0.15 for intersubject decoding, as the training set is much larger).

The two pooling operations yield a cumulative temporal reduction factor of 16, corresponding to an effective temporal resolution of approximately 32 ms. This scale preserves relevant EEG temporal structure while limiting feature dimensionality and reducing overfitting risk.

#### Fully connected and output layers

Feature maps are flattened and passed to a fully connected layer with two output units corresponding to the decoding classes. A max-norm constraint (norm rate = 0.25) is applied to improve generalization. The output layer applies a softmax activation function to produce normalized class probability estimates. Final predictions correspond to the class with highest posterior probability.

#### Activation functions and hyperparameter selection

ELU nonlinearities are applied after batch normalization in all convolutional layers to model non-linear EEG patterns while preserving stable gradient propagation. The output layer uses softmax activation.

Hyperparameters governing model capacity, including filter counts, depth multiplier, and dropout rates, were tuned using a preliminary grid search and fixed for all final decoding analyses. Temporal-scale parameters (kernel length and pooling factors) were instead fixed a priori based on sampling characteristics and physiologically relevant time scales, as they determine temporal feature granularity rather than model flexibility.

### 3.2 Input representation and data preparation

EEG epochs obtained after preprocessing (Section 2.7) served as network inputs. Epoch lengths were adapted to each experiment while remaining strictly confined to the pre-feedback interval.

Data were organized as arrays of shape (N × channels × samples), where N denotes trial count. Experiment 1 epochs contained ∼280 samples (∼560 ms at 500 Hz), whereas Experiment 2 epochs contained ∼420 samples (∼840 ms).

All data were converted to single-precision floating-point format. Non-finite values were replaced with zeros. These procedures were applied identically across training, validation, and test datasets and did not involve label-dependent processing.

### 3.3 Feature standardization and tensor construction

Z-score normalization was applied independently for each EEG channel. Training data were reshaped to (N × samples, channels) to estimate channel-wise mean and standard deviation. A scaler fitted exclusively on training data was applied unchanged to validation and test sets, preventing data leakage.

After normalization, tensors were reshaped back to trial format and expanded with a singleton dimension. Final input shapes were (N, 17, 280, 1) for Experiment 1 and (N, 17, 420, 1) for Experiment 2. Labels were integer-encoded and converted to one-hot vectors (N × 2).

### 3.4 Model training and optimization

Models were implemented in TensorFlow/Keras with identical optimization settings across experiments. Training used the Adam optimizer with initial learning rate η (0.0006 for Experiment 1 and 0.001 for Experiment 2; see Supplementary Results section, Tables 3 & 7) and categorical cross-entropy loss. Models were trained for a maximum of 200 epochs with a batch size of 16. Training was performed on a single GPU.

Model hyperparameters, including filter counts, depth multiplier, dropout rate, learning rate and batch size, were determined through a preliminary grid search over predefined ranges based on validation performance (see Supplementary Results, Tables 3-4 & 7-8). Hyperparameter optimization was completed before final analyses, and the selected configuration was fixed thereafter.

Validation data were used exclusively for convergence monitoring and regularization after hyperparameter selection. Training stability was ensured using two callbacks monitored on validation loss: learning rate scheduler (ReduceLROnPlateau with factor = 0.8, patience = 8, minimum learning rate = 1e-6) and early stopping (patience = 20, minimum = 1e-3), restoring weights from the minimum validation loss.

Data partitioning into training, validation, and test sets was performed at the block level for intra-subject decoding and at the participant level for inter-subject decoding (Section 3.5). The test set was strictly held out from all training and optimization stages.

Model performance was evaluated on held-out data using accuracy, precision, recall, F1-score, and AUC.

### 3.5 Decoding strategy and evaluation protocol

#### Intra-subject decoding

In the intra-subject setting, decoding models were trained and evaluated independently for each participant, yielding participant-specific classifiers. No data were pooled across participants at any stage of the analysis.

To preserve the temporal organization of the recording session and minimize information leakage arising from temporal dependencies in EEG activity, data were partitioned at the block level rather than at the trial level. For each participant and each experimental condition, the dataset comprised eight sequentially acquired blocks, which served as the elementary units for data splitting.

For a given partition, blocks were divided into mutually exclusive training, validation, and test sets according to a 4/2/2 scheme. Block assignments were constrained such that the three subsets jointly sampled early, intermediate, and late phases of the session. This constraint was introduced to reduce potential temporal confounds, including practice effects, fatigue, and slow non-stationarities in the EEG signal.Within a given partition, identical block indices were used for the two classes entering the binary classifier (e.g., human-controlled versus AI-controlled interaction in Experiment 1; AI with explanation versus AI without explanation in Experiment 2). This ensured that class discrimination could not be attributed to block-specific or session-specific fluctuations unrelated to the experimental manipulation.

To obtain stable performance estimates, this partitioning procedure was repeated across five distinct block configurations consistent with the 4/2/2 split. Across repetitions, different combinations of blocks were assigned to the validation and test sets in order to reduce the likelihood that decoding performance would be biased by a single block assignment and therefore depend idiosyncratically on a specific choice of blocks. Performance metrics were computed independently for each repetition and then averaged to obtain the final intra-subject estimate for each participant.

Validation data were used exclusively for training-control procedures, including early stopping and adaptive learning-rate scheduling. Test data remained strictly held out throughout model fitting, model selection, and regularization, and were used only for final performance evaluation.

#### Inter-subject decoding

Inter-subject generalization was assessed using a leave-one-subject-out (LOSO) cross-validation procedure. Whereas intra-subject decoding tested whether agency-related neural states could be detected reliably within individual participants, inter-subject decoding assessed whether these states generalized across participants. This distinction is important for neuroadaptive applications, as cross-participant generalization would support the development of systems that do not require complete recalibration for each new user.

**Table 2.**
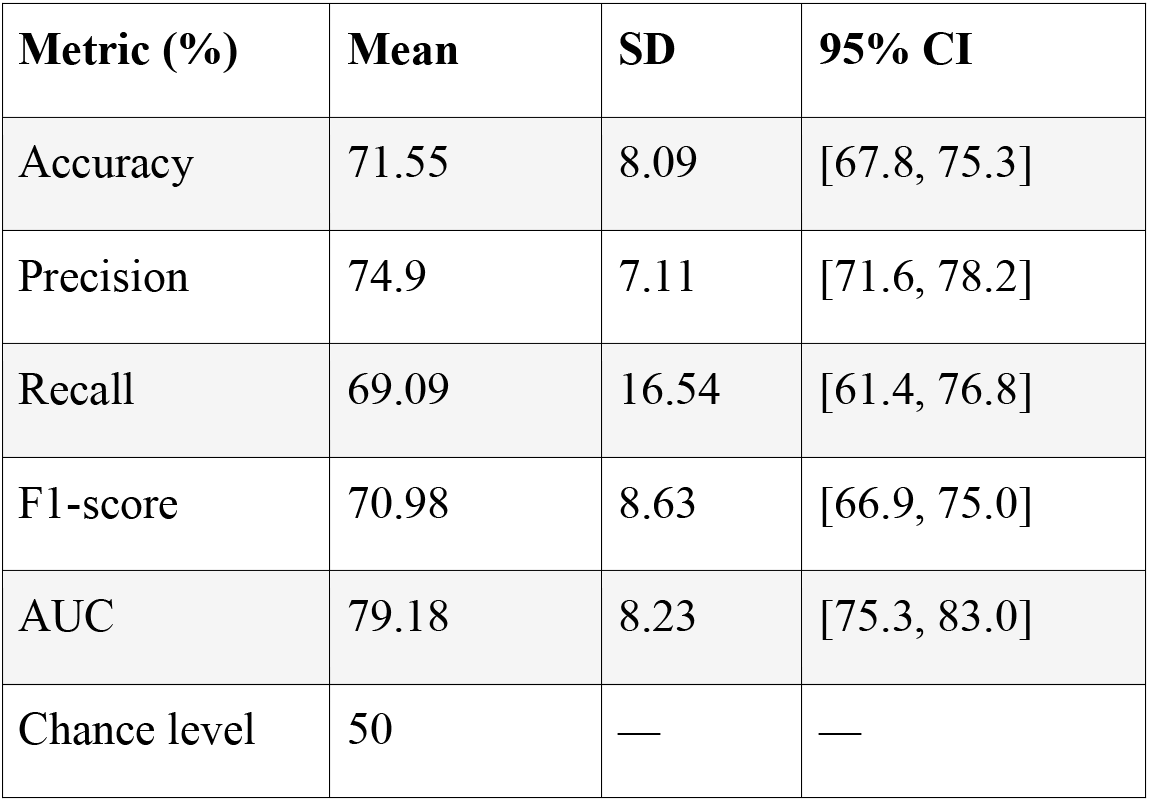
Inter-subject decoding performance in Experiment 1 under a leave-one-subject-out (LOSO) evaluation scheme (N=20). Accuracy, Precision, Recall, F1-score, and AUC were computed for each held-out participant on strictly unseen test data and averaged across five runs using different validation participants to obtain a single participant-level estimate. Group-level means, standard deviations, and 95% t-based confidence intervals were then computed across participants. The chance level for binary classification was 50%.

| Metric (%) | Mean | SD | 95% CI |
| --- | --- | --- | --- |
| Accuracy | 71.55 | 8.09 | [67.8, 75.3] |
| Precision | 74.9 | 7.11 | [71.6, 78.2] |
| Recall | 69.09 | 16.54 | [61.4, 76.8] |
| F1-score | 70.98 | 8.63 | [66.9, 75.0] |
| AUC | 79.18 | 8.23 | [75.3, 83.0] |
| Chance level | 50 | — | — |

In each LOSO iteration, one participant was held out entirely as the test set, and all remaining participants except one were used for training. To reduce dependence on the identity of the validation participant, each LOSO iteration was repeated across five folds while keeping the same held-out test participant. In each fold, one validation participant was selected at random from the remaining available participants, and all other participants were assigned to the training set. Thus, for a given test subject, model training and training-control procedures were evaluated against five distinct validation subjects. This procedure reduced the likelihood that inter-subject decoding performance would depend idiosyncratically on the choice of validation participant.

Training, validation, and test sets were therefore strictly separated at the participant level, preventing any leakage of subject-specific information across partitions. As in the intra-subject setting, validation data were used exclusively for training-control procedures, whereas the held-out test participant was used only for final model evaluation. The LOSO procedure was repeated until each participant had served once as the test subject. Performance metrics were computed separately for each fold and then averaged across folds and test participants to obtain the final estimate of inter-subject decoding performance.

#### Class balance

Class distributions were balanced by design in both intra-subject and inter-subject decoding analyses. Accordingly, no additional class weighting, resampling, or cost-sensitive learning procedures were applied during model training.

### 3.6 Evaluation Metrics

Model performance was assessed using multiple complementary evaluation metrics to provide a comprehensive characterization of decoding performance and to facilitate comparison with prior EEG decoding studies. Specifically, we report categorical accuracy, precision, recall, and the F1-score.

Accuracy was used as a global measure of classification performance, quantifying the proportion of correctly classified trials across all classes. Accuracy is defined as:

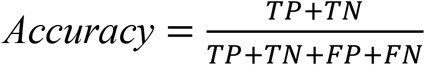

where *TP*,*TN*,*FP*,*FN* denote the numbers of true positives, true negatives, false positives, and false negatives, respectively.

Although accuracy provides an overall summary of model performance, it can be misleading in the presence of class imbalance or asymmetric classification errors. To address this limitation, precision and recall were additionally computed to quantify class-specific performance. Precision reflects the proportion of predicted positive instances that are correctly classified, whereas recall (also referred to as sensitivity) measures the proportion of true positive instances that are correctly detected. These metrics are defined as:

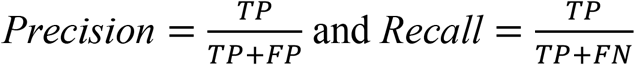

To further capture the balance between precision and recall, the F1-score was computed as their harmonic mean:

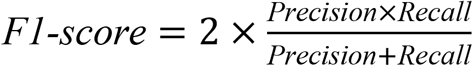

The F1-score provides a robust summary of classification performance by penalizing extreme imbalances between precision and recall and is particularly informative when evaluating models under conditions where class-wise performance is critical.

In addition to these threshold-dependent metrics, we computed the area under the receiver operating characteristic curve (AUC). The ROC curve characterizes the trade-off between the true positive rate and the false positive rate across all possible decision thresholds. The AUC therefore provides a threshold-independent measure of class separability and reflects the probability that the classifier assigns a higher score to a randomly chosen positive instance than to a randomly chosen negative instance. In binary classification, an AUC of 50% corresponds to chance-level separability, whereas a value of 100% indicates perfect discrimination between classes.

All metrics were computed exclusively on held-out test data and were aggregated consistently with the evaluation procedures described above. For intra-subject decoding, metrics were first computed on each held-out cross-validation fold and then averaged across folds within each participant. For inter-subject decoding, metrics were computed on the held-out participant in each leave-one-subject-out iteration and averaged across the five validation runs. These procedures yielded stable participant-level estimates that were subsequently used for group-level statistical analyses.

### 3.7 Frequency-band ablation analysis

To investigate the contribution of canonical EEG frequency bands to decoding performance, a frequency-band ablation analysis was conducted. EEG signals were decomposed into standard frequency ranges: delta (2–4 Hz), theta (4–8 Hz), alpha (8–13 Hz), and beta (13–30 Hz), consistent with the frequency ranges used in the time–frequency analysis (Section 4).

Band-specific ablation was performed by selectively removing one frequency band at a time using zero-phase band-stop filtering, thereby preserving the remaining spectral content while eliminating the targeted frequency component. Filtering was applied after preprocessing and prior to feature standardization, ensuring that all subsequent processing steps remained identical across conditions.

For each ablation condition, the modified EEG signals were processed using the same pipeline described in Sections 3.2–3.4. The EEGNet architecture, training procedure, and all hyperparameters were kept strictly identical to the baseline configuration. The only difference between conditions was the removal of a specific frequency band from the input.

Decoding performance under each ablation condition was evaluated using the same procedures described above. For the ablation analysis, participant-level performance scores were obtained by averaging test metrics across all evaluation folds within each participant, such that the participant constituted the unit of statistical inference.

Statistical comparisons were performed between each ablation condition and the baseline (full-band) condition using paired two-sided t-tests across participants. This analysis was conducted separately for each decoding metric of interest, including area under the receiver operating characteristic curve (AUC) and classification accuracy when available.

To control for multiple comparisons across frequency bands and performance metrics, p-values were adjusted using the Benjamini–Hochberg false discovery rate (FDR) procedure. Statistical significance was assessed at a corrected threshold of α = 0.05. In addition to p-values, effect sizes were quantified using Cohen’s d, computed from within-participant differences between baseline and ablation conditions.

Ablation analyses were performed independently for intra-subject and inter-subject decoding settings, following the same data partitioning and evaluation protocols described above. Accordingly, observed performance differences are interpretable as reflecting the contribution of frequency-specific information rather than differences in model architecture, training procedure, or evaluation strategy.

## 4. Time-Frequency Analysis

Time–frequency analyses (TFRs) were conducted to characterize condition-dependent oscillatory dynamics at the group level and were not intended to support single-trial classification. Accordingly, TFR results are interpreted as complementary, descriptive markers of neural processes underlying the decoding results. TFRs of EEG power were computed using the multitaper method as implemented in MNE-Python. Spectral power was estimated between 2 and 30 Hz using 29 linearly spaced frequency bins, covering the canonical delta (2–4 Hz), theta (4–8 Hz), alpha (8–13 Hz), and beta (13–30 Hz) ranges. Frequency-dependent temporal windows were used, with the number of cycles defined as half the frequency value. Spectral smoothing was controlled using a time–bandwidth product of 3.0, corresponding to 5 discrete prolate spheroidal sequence (DPSS) tapers.

TFRs were computed separately for each participant and condition using extended peri-action epochs in order to ensure sufficient temporal context for baseline normalization and stable estimation of low-frequency power. The duration of these visualization epochs differed across experiments because the available pre-keypress interval was constrained by task timing. In Experiment 1, pre-keypress periods were relatively short because their duration depended solely on participants ‘reaction times; accordingly, epochs used for visualization spanned from −700 to +700 ms relative to keypress. In Experiment 2, longer pre-keypress intervals were available because reaction times were preceded by an additional mandatory 600-ms explanation display period; visualization epochs therefore extended from −1200 to +1200 ms relative to keypress.

For each condition, power estimates were averaged across trials during TFR computation, yielding one condition-specific TFR per participant. Inter-trial coherence was not computed. Baseline correction was then applied using a log-ratio transform. In Experiment 1, the baseline interval was defined as −500 to −300 ms relative to keypress, whereas in Experiment 2 the baseline interval was defined as −900 to −700 ms. Baseline-corrected power was computed as:

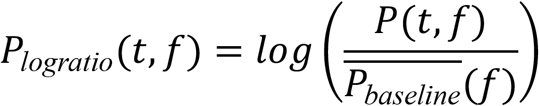

where *P*(*t*, *f*) denotes the spectral power at time *t* and frequency *f*, and 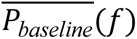 represents the mean power computed within the baseline interval for frequency *f*.

Although TFRs were computed on extended epochs, statistical analyses were restricted to the peri-action windows used in the decoding analyses in order to ensure direct correspondence with the EEGNet input segments. Accordingly, band-limited power was extracted from −280 to +280 ms relative to keypress in Experiment 1 and from −560 to +280 ms in Experiment 2. Power values were then averaged within canonical frequency bands and entered into participant-level statistical comparisons between conditions.

Condition differences were assessed using paired two-sided t-tests at the participant level. To control for multiple comparisons across frequency bands, p-values were adjusted using the Benjamini–Hochberg false discovery rate (FDR) procedure.

## 5. Results

### 5.1. Behavioral validation of agency manipulation

To verify that the experimental manipulation effectively modulated participants’ subjective sense of agency, self-reported Feeling of Control (FoC) ratings were collected on 20% of trials using a 9-point scale. These ratings provided a behavioral validation of the agency manipulation prior to the neural decoding analyses.

#### 5.1.1 Experiment 1: Effect of decision authority on FoC ratings

Delegating decision authority to the AI significantly reduced perceived control, F (1, 31.4) = 16.25, p = 0.00033, η²ₚ = 0.94 (see Supplementary Results section-Chapter III-Figure 1A). Tukey-corrected post hoc comparisons revealed that FoC ratings were higher in the Motor condition (M = 7.52, SD = 0.15, 95% CI [7.22, 7.82]) than in the AI condition (M = 6.80, SD = 0.26, 95% CI [6.27, 7.33]; β = 0.721, t (31.8) = 3.57, p = 0.0012).

These findings confirm that the manipulation of decision authority reliably altered participants ‘ explicit sense of control.

#### 5.1.2 Experiment 2: Effect of AI explanatory information on FoC ratings

Providing distal, goal-level explanations significantly increased perceived control relative to the no-explanation condition, F (1, 31.5) = 15.83, p = 0.00038, η²ₚ = 0.96 (see Supplementary Results section-Chapter III-Figure 1B). Tukey-corrected comparisons showed that FoC ratings were lower in the No-explanation condition (M = 6.08, SE = 0.28, 95% CI [5.50, 6.65]) than when Distal explanations were provided (M = 6.81, SE = 0.22, 95% CI [6.36, 7.26]). This effect was further supported by the mixed-effects model (β = −0.735, t(32.5) = −3.91, p = 0.0006).

Together, these results demonstrate that system explainability significantly enhances subjective control under automated decision-making.

Crucially, across both experiments, subjective ratings confirmed that decision authority and explanatory context systematically modulated perceived agency. These behavioral effects establish that the task successfully induced meaningful variations in agency, thereby providing a principled basis for testing whether such variations can be decoded from pre-feedback EEG activity using deep learning approaches.

### 5.2. Decoding Agency performance evaluation

Decoding performance was quantified at the participant level using complementary metrics computed exclusively on held-out test data. Accuracy was used as the primary measure of classification performance, and was complemented by precision, recall, F1-score, and the area under the receiver operating characteristic curve (AUC). These additional metrics were included to verify that decoding performance was not driven by asymmetric class predictions or by decision-threshold-dependent bias.

In the intra-subject setting, models were trained and evaluated independently for each participant using five repeated block-wise train/validation/test partitions. For each repetition, data were split according to a 4/2/2 block scheme, with different block assignments across repetitions to reduce the dependence of performance estimates on a single validation/test split. For each repetition, accuracy, precision, recall, F1-score, and AUC were computed on the held-out test set. Metrics were then averaged across repetitions to obtain one estimate per participant. Group-level statistics were subsequently computed across participants (N = 20 for Experiment 1 and N = 17 for Experiment 2).

In the inter-subject setting, cross-participant generalisation was assessed using a leave-one-subject-out (LOSO) procedure. At each iteration, one participant was held out entirely as the test set, one additional participant was used as the validation set, and all remaining participants were assigned to the training set. To reduce dependence on the identity of the validation participant, each LOSO iteration was repeated five times while keeping the same held-out test participant and varying the validation participant across runs. Accuracy, precision, recall, F1-score, and AUC were computed on the held-out test participant for each run, then averaged across the five runs to obtain one estimate per held-out participant. Final inter-subject performance was summarised across held-out participants.

This procedure ensured that all reported metrics were derived exclusively from unseen test data and that performance estimates were not driven by a specific block partition, validation block, or validation participant.

#### 5.2.1 Experiment 1: Decoding of the Control Delegation Effect

##### Intra-subject decoding

In Experiment 1, intra-subject decoding assessed whether pre-feedback EEG activity discriminated between Motor/no-AI and belief-based AI trials, corresponding to different levels of control delegation.

**Figure 2.**
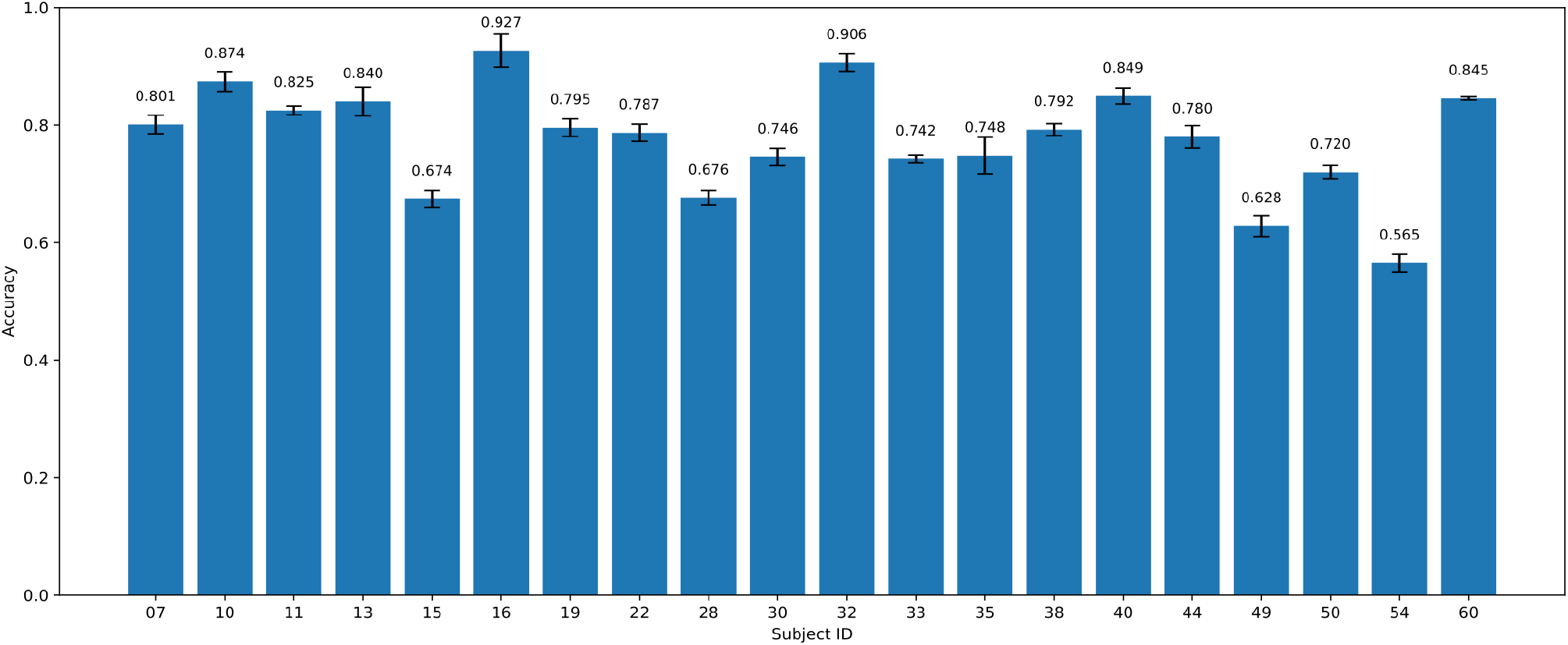
Intra-subject decoding performance in Experiment 1 (N=20). Each bar represents the mean test accuracy obtained for an individual participant, computed on strictly held-out data and averaged across five repeated block-wise train/validation/test partitions. Error bars indicate the standard deviation across repetitions. Numerical values above the bars denote participant-level mean accuracies. Across participants, mean cross-validated test accuracy reached 77.61% (SD = 9.17%; 95% CI: [73.30, 81.90]), well above the theoretical chance level of 50% for binary classification.

Accuracy was also stable across folds within participants, with a mean within-participant standard deviation of 3.90%. A two-sided one-sample t-test confirmed that accuracy was significantly above chance, t(19) = 13.46, p < 1 × 10⁻¹⁰, with a very large effect size (Cohen’s d = 3.01).

Complementary metrics showed the same pattern. Mean F1-score reached 77.42% (SD = 9.36%; 95% CI: [73.04, 81.80]), mean precision reached 78.84% (SD = 9.05%; 95% CI: [74.60, 83.08]), and mean recall reached 78.87% (SD = 8.92%; 95% CI: [74.69, 83.04]). These values indicate balanced classification performance across class labels. Mean AUC reached 84.25% (SD = 9.73%; 95% CI: [79.70, 88.80]), indicating robust class separability independently of a fixed decision threshold.

Statistical tests against chance confirmed that all complementary metrics were significantly above 50%: F1-score, t(19) = 14.88, p < 1 × 10⁻¹⁰; precision, t(19) = 14.24, p < 1 × 10⁻¹¹; recall, t(19) = 14.46, p < 1 × 10⁻¹¹; and AUC, t(19) = 15.74, p < 1 × 10⁻¹². All effects were very large, with Cohen’s d values greater than 2.90.

Together, these results indicate that pre-feedback EEG activity contained reliable information about the level of control delegation at the single-participant level, and that decoding performance reflected discriminative neural structure rather than class imbalance, response bias, or threshold-dependent effects.

##### Inter-subject decoding

Inter-subject decoding assessed whether neural patterns related to control delegation generalised across participants. Across held-out participants, mean decoding accuracy reached 71.55% (SD = 8.09%; 95% CI: [67.77, 75.34]), with individual accuracies ranging from 56.20% to 85.62%. A two-sided one-sample t-test against chance confirmed that accuracy was significantly above 50%, t(19) = 11.92, p = 2.92 × 10⁻¹⁰, with a very large effect size (Cohen’s d = 2.66).

Complementary metrics supported this result. Mean precision reached 74.90% (SD = 7.11%; 95% CI: [71.60, 78.20]), mean recall reached 69.09% (SD = 16.54%; 95% CI: [61.40, 76.80]), and the corresponding F1-score was 70.98% (SD = 8.63%; 95% CI: [66.90, 75.00]). Mean AUC reached 79.18% (SD = 8.23%; 95% CI: [75.30, 83.00]), indicating robust class separability independently of a fixed decision threshold.

Statistical inference confirmed that all complementary metrics exceeded chance level: precision, t(19) = 15.70, p = 3.20 × 10⁻¹², Cohen’s d = 3.51; recall, t(19) = 5.16, p = 5.60 × 10⁻⁵, Cohen’s d = 1.15; F1-score, t(19) = 10.90, p = 3.60 × 10⁻⁹, Cohen’s d = 2.44; and AUC, t(19) = 15.85, p = 2.60 × 10⁻¹², Cohen’s d = 3.54.

Although inter-subject decoding is inherently more demanding than intra-subject decoding because of inter-individual variability in neural dynamics, the model maintained robust above-chance performance under strict participant-wise separation. Recall showed greater variability than precision, suggesting some inter-individual differences in class-specific sensitivity. Nevertheless, the convergence of accuracy, precision, recall, F1-score, and AUC indicates that inter-subject decoding performance was not driven by asymmetric predictions, class imbalance, or decision-threshold bias. These results suggest that the discriminative EEG patterns captured by the model reflected neural representations that generalized across participants rather than purely subject-specific idiosyncrasies.

#### 5.2.2 Experiment 2: Decoding of AI Explanatory Information

##### Intra-subject decoding

In Experiment 2, intra-subject decoding was used to assess whether pre-feedback EEG activity discriminated between belief-based AI trials with and without distal explanatory information. Across participants, mean cross-validated test accuracy reached 62.50% (SD = 11.98%; 95% CI: [56.34, 68.66]), with a mean within-participant standard deviation across folds of 3.84%.

**Figure 3.**
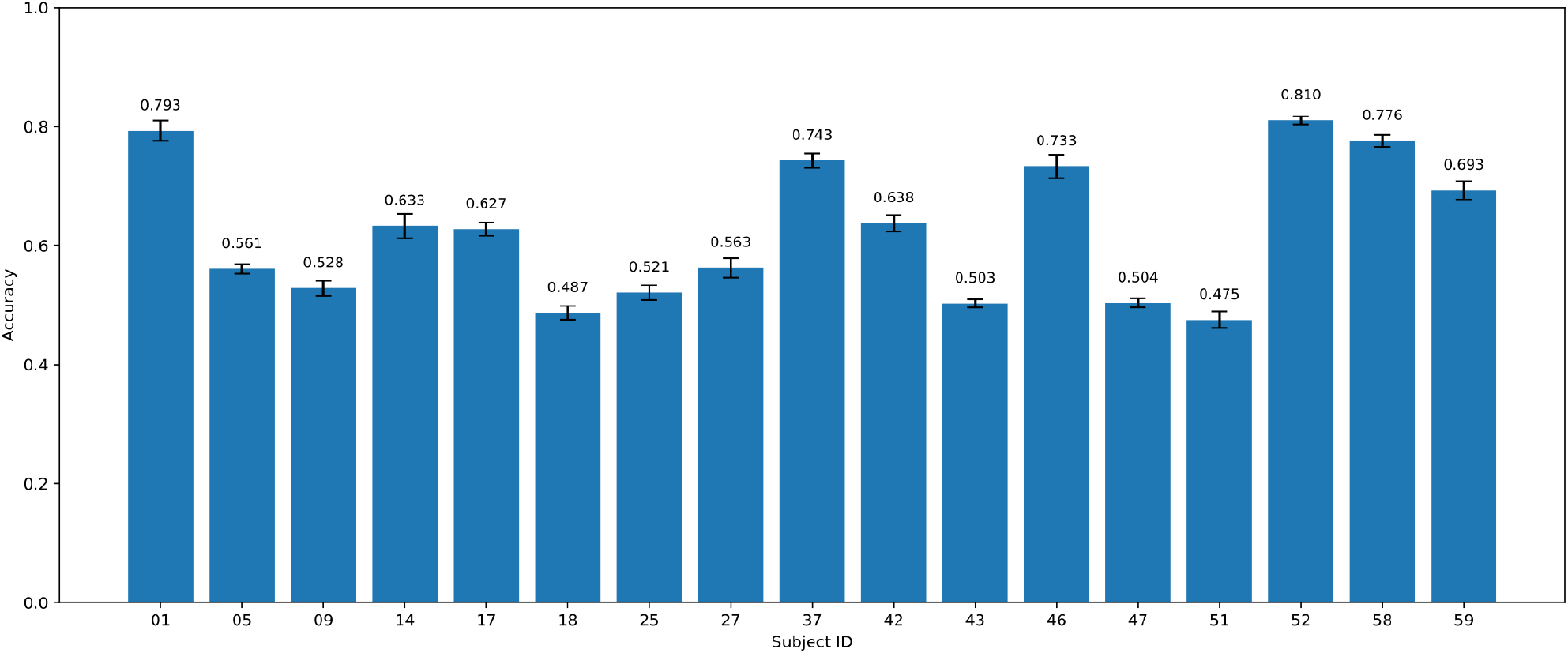
Intra-subject decoding performance in Experiment 2 (N=17). Each bar represents the mean test accuracy obtained for an individual participant, computed on strictly held-out data and averaged across five repeated block-wise train/validation/test partitions. Error bars indicate the standard deviation across repetitions. Numerical values above the bars denote participant-level mean accuracies.

A two-sided one-sample t-test against the theoretical chance level of 50% confirmed that accuracy was significantly above chance, t(16) = 4.30, p = 5.49 × 10⁻⁴, with a large effect size (Cohen’s d = 1.04).

Complementary metrics converged with the accuracy results. Mean precision reached 62.74% (SD = 13.02%; 95% CI: [56.30, 69.20]), mean recall reached 66.05% (SD = 10.91%; 95% CI: [60.60, 71.50]), and the corresponding F1-score was 63.28% (SD = 10.92%; 95% CI: [57.90, 68.70]). The close correspondence between precision and recall indicates that decoding performance was not driven by a systematic bias toward one class. Mean AUC reached 66.11% (SD = 15.19%; 95% CI: [58.60, 73.70]), indicating reliable, albeit moderate, class separability independently of a fixed decision threshold.

Statistical tests against chance confirmed that all complementary metrics exceeded 50%: precision, t(16) = 4.03, p = 9.60 × 10⁻⁴, Cohen’s d = 0.98; recall, t(16) = 6.07, p = 1.64 × 10⁻⁵, Cohen’s d = 1.47; F1-score, t(16) = 5.01, p = 1.27 × 10⁻⁴, Cohen’s d = 1.22; and AUC, t(16) = 4.37, p = 4.73 × 10⁻⁴, Cohen’s d = 1.06.

**Table 3.** Intra-subject decoding performance in Experiment 2 (N=17). Accuracy, Precision, Recall, F1-score, and AUC were computed for each participant on strictly held-out test data and averaged across five repeated block-wise train/validation/test partitions to obtain a single participant-level estimate. Group-level means, standard deviations, and 95% t-based confidence intervals were then computed across participants. Chance level for binary classification was 50%.

| Metric (%) | Mean | SD | 95% CI |
| --- | --- | --- | --- |
| Accuracy | 62.5 | 11.98 | [56.5, 68.5] |
| Precision | 62.74 | 13.02 | [56.3, 69.2] |
| Recall | 66.05 | 10.91 | [60.6, 71.5] |
| F1-score | 63.28 | 10.92 | [57.9, 68.7] |
| AUC | 66.11 | 15.19 | [58.6, 73.7] |
| Chance level | 50 | — | — |

Together, these results indicate that pre-feedback EEG activity encoded information about the availability of distal explanatory information during belief-based AI decision-making. Although decoding performance was more moderate than in Experiment 1, the convergence of accuracy, precision, recall, F1-score, and AUC suggests that classification reflected genuine discriminative neural information rather than response bias, class imbalance, or threshold-dependent effects.

##### Inter-subject decoding

Inter-subject decoding was used to assess whether neural patterns related to AI explanatory information generalized across participants. Across held-out participants, mean decoding accuracy reached 58.90% (SD = 8.20%; 95% CI: [54.70, 63.10]), with individual accuracies ranging from 47.80% to 73.10%.

A two-sided one-sample t-test against the theoretical chance level of 50% confirmed that accuracy was significantly above chance, t(16) = 4.48, p = 3.78 × 10⁻⁴, with a large effect size (Cohen’s d = 1.09).

**Table 4.**
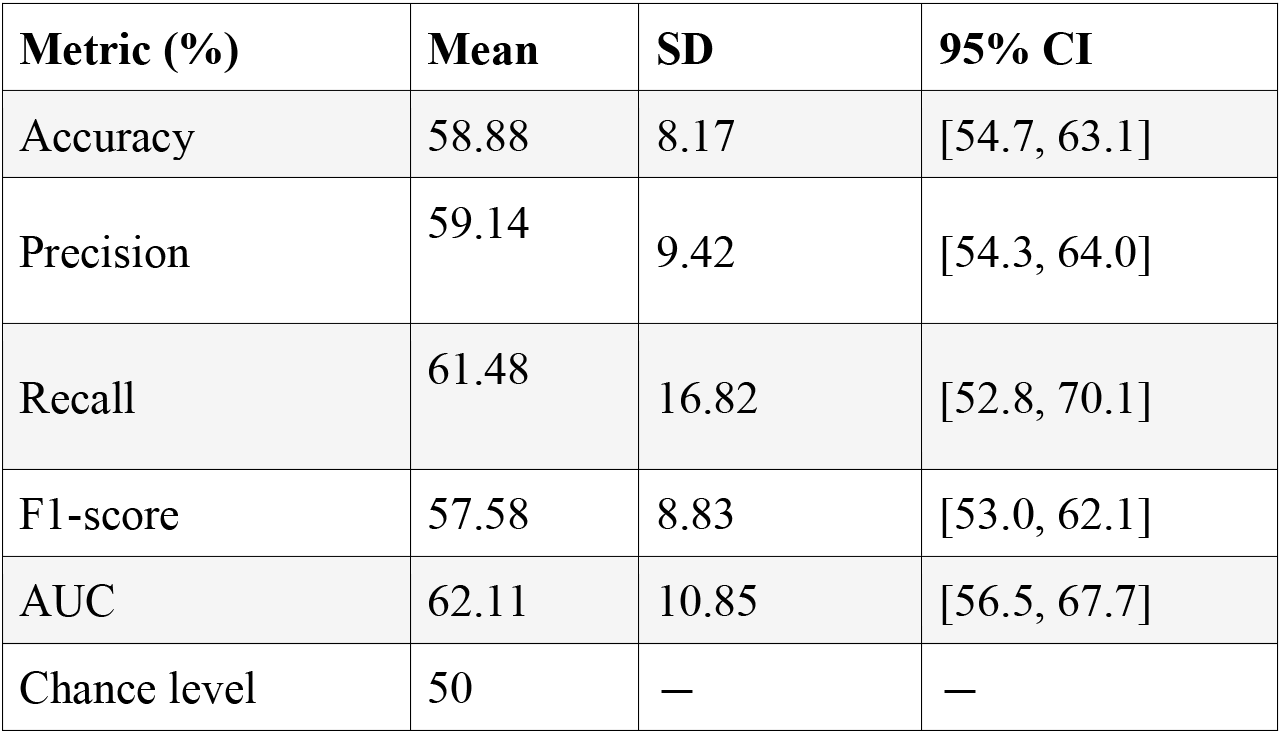
Cross-validated inter-subject decoding performance in Experiment 2. Accuracy, Precision, Recall, F1-score, and AUC were computed on held-out test participants under the leave-one-subject-out evaluation scheme and averaged across decoding runs to obtain participant-level performance estimates. Group-level means, standard deviations, and 95% t-based confidence intervals were then calculated across held-out participants. Chance level for binary classification was 50%.

| **Metric (%)** | **Mean** | **SD** | **95% CI** |
| --- | --- | --- | --- |
| Accuracy | 58.88 | 8.17 | [54.7, 63.1] |
| Precision | 59.14 | 9.42 | [54.3, 64.0] |
| Recall | 61.48 | 16.82 | [52.8, 70.1] |
| F1-score | 57.58 | 8.83 | [53.0, 62.1] |
| AUC | 62.11 | 10.85 | [56.5, 67.7] |
| Chance level | 50 | — | — |

Complementary metrics showed a consistent pattern. Mean precision reached 59.14% (SD = 9.42%; 95% CI: [54.30, 64.00]), mean recall reached 61.48% (SD = 16.82%; 95% CI: [52.80, 70.10]), and the corresponding F1-score was 57.58% (SD = 8.83%; 95% CI: [53.00, 62.10]). The broad similarity between precision and recall suggests that decoding was not driven by strongly asymmetric class predictions, although the higher variability in recall indicates some inter-individual differences in class-specific sensitivity. Mean AUC reached 62.11% (SD = 10.85%; 95% CI: [56.50, 67.70]), supporting the presence of discriminative information independently of a fixed decision threshold.

Statistical tests against chance confirmed that all complementary metrics exceeded 50%: precision, t(16) = 4.00, p = 1.03 × 10⁻³, Cohen’s d = 0.97; recall, t(16) = 2.81, p = 0.0125, Cohen’s d = 0.68; F1-score, t(16) = 3.54, p = 2.73 × 10⁻³, Cohen’s d = 0.86; and AUC, t(16) = 4.60, p = 2.95 × 10⁻⁴, Cohen’s d = 1.12.

Although inter-subject decoding was weaker than intra-subject decoding, the classifier remained reliably above chance under strict participant-wise separation.

Together, the convergence of accuracy, precision, recall, F1-score, and AUC indicates that inter-subject decoding performance reflected genuine class-discriminative structure in the EEG signal rather than trivial response bias, class imbalance, or threshold-dependent effects. These findings suggest that explanatory information in belief-based AI decision-making was associated with neural patterns that generalized across individuals, albeit with moderate inter-subject variability.

### 5.3 Frequency-Band Contribution to Decoding Performance: the oscillatory ablation effect

To quantify the spectral contributions underlying decoding performance, we conducted systematic frequency-band ablation analyses in both experiments. For each condition, models were retrained after removing one canonical EEG frequency band at a time (delta, theta, alpha, beta), while keeping all preprocessing, architectural, and training parameters identical to the baseline model.

Performance metrics were computed at the participant level using the same evaluation procedures described in Section 3.6. For intra-subject decoding, models were evaluated using block-based cross-validation within each participant, and accuracy and AUC were averaged across folds to obtain a single participant-level estimate. For inter-subject decoding, generalization was assessed using a leave-one-subject-out (LOSO) scheme, where performance for each held-out participant was obtained by averaging results across the five validation runs.

Group-level statistics were computed across participant-level estimates. Differences between the full-spectrum baseline and each ablated condition were evaluated using two-tailed paired t-tests, with Benjamini–Hochberg false discovery rate (FDR) correction applied across band contrasts and metrics.

#### 5.4.1 Experiment 1: Oscillatory ablation impact on decoding Decision Authority

##### Intra-subject

**Baseline** performance reached 77.61% of accuracy and AUC = 84.25% at the participant level. Removing the **delta band** produced the largest decrement in decoding performance, reducing accuracy to 71.30% and AUC to 77.31% (ΔAccuracy = −6.31 pp; ΔAUC = −6.94 relative to baseline). This reduction was significant for both metrics after FDR correction (Accuracy: t(19) = 4.235, p = 0.0042; AUC: t(19) = 3.778, p = 0.0042), indicating that delta-band activity carried a substantial proportion of the discriminative information.

Removal of the **theta band** yielded smaller but still reliable reductions in performance. Accuracy decreased to 75.78% and AUC to 82.64% (ΔAccuracy = −1.83 pp; ΔAUC = −1.61 relative to baseline). These effects also remained significant after correction (Accuracy: t(19) = 2.935, p = 0.0121; AUC: t(19) = 2.598, p = 0.0196), suggesting a secondary contribution of theta-band activity to decoding performance.

**Table 5.** Experiment 1: Spectral Ablation Analysis (Intra-subject decoding). Mean decoding performance across participants after removal of canonical EEG frequency bands (delta, theta, alpha, beta). Values indicate mean ±standard deviation across participants, where participant-level estimates were obtained by averaging test performance across cross-validation folds within each participant. Baseline corresponds to the full-spectrum model, and band ablations were implemented by retraining the model after removing the corresponding frequency band from the input signal. The largest performance reduction is observed following delta-band ablation, indicating that slow-frequency activity provides the strongest contribution to decoding decision authority.

| Condition | Accuracy (%) | AUC (%) |
| --- | --- | --- |
| Baseline | 77.61 ±9.17 | 84.25 ±9.73 |
| – Delta | 71.30 ±11.74 | 77.31 ±13.49 |
| – Theta | 75.78 ±8.40 | 82.64 ±9.79 |
| – Alpha | 76.88 ±8.88 | 83.81 ±9.74 |
| – Beta | 77.63 ±8.88 | 84.46 ±9.44 |

In contrast, removing the **alpha band** produced only a negligible reduction in performance, with accuracy reaching 76.88% and AUC 83.81% (ΔAccuracy = −0.73 pp; ΔAUC = −0.44 relative to baseline). These differences were not statistically significant after FDR correction (Accuracy: t(19) = 1.726, p = 0.1366; AUC: t(19) = 1.136, p = 0.2893), indicating that alpha-band activity did not contribute substantially to decoding performance in the present setting.

Finally, removing the **beta band** had virtually no effect on decoding performance. Accuracy remained unchanged at 77.63%, and AUC showed only a minimal increase to 84.46% (ΔAccuracy = +0.02 pp; ΔAUC = +0.21 relative to baseline). These differences were not statistically significant after FDR correction (Accuracy: t(19) = −0.071, p = 0.9439; AUC: t(19) = −0.872, p = 0.5253), indicating that beta-band activity did not make a reliable contribution to decoding performance in the present setting.

Overall, these ablations support a dominant contribution of low-frequency components (delta, then theta) to decoding decision authority, with limited or no evidence that alpha/beta bands are necessary for robust performance under the present pipeline.

##### Inter-Subject

Baseline performance under the LOSO framework reached a mean accuracy of 71.55 % and an AUC of 79.18% across participants.

**Table 6.** Inter-subject ablation analysis in Experiment 1 under a leave-one-subject-out (LOSO) evaluation scheme(N=20). Mean decoding Accuracy and AUC are reported for the baseline model and for band-ablation models in which one canonical frequency band (delta, theta, alpha, or beta) was removed at a time. Performance was computed on strictly held-out test participants and averaged across five runs using different validation participants to obtain participant-level estimates. Values are expressed as mean ±standard deviation across participants.

| Condition | Accuracy (%) | AUC (%) |
| --- | --- | --- |
| Baseline | 71.55 ±8.09 | 79.18 ±8.23 |
| – Delta | 68.48 ±8.90 | 75.73 ±9.11 |
| – Theta | 71.53 ±7.09 | 79.30 ±7.58 |
| – Alpha | 71.52 ±7.68 | 79.18 ±8.25 |
| – Beta | 71.30 ±7.73 | 79.03 ±8.14 |

Removing the **delta band** produced the largest degradation in decoding performance. Accuracy decreased to 68.48% and AUC to 75.73% (ΔAccuracy = −3.07 pp; ΔAUC = −3.45 relative to baseline). Paired t-tests confirmed that these reductions were statistically significant after FDR correction (Accuracy: t(19) = 3.359, p = 0.0132; AUC: t(19) = 3.390, p = 0.0132), indicating that delta-band activity contributes substantially to neural representations that generalize across participants.

**Table 7.** Experiment 2: Spectral Ablation Analysis (Intra-subject decoding). Mean decoding Accuracy and AUC are reported for the baseline model and for band-ablation models in which one canonical frequency band (delta, theta, alpha, or beta) was removed at a time. Performance was computed for each participant on strictly held-out test data and averaged across five repeated block-wise train/validation/test partitions to obtain participant-level estimates. Values are expressed as mean ±standard deviation across participants.

| Condition | Accuracy (%) | AUC (%) |
| --- | --- | --- |
| Baseline | 62.50 ±11.98 | 66.11 ±15.19 |
| – Delta | 57.50 ±11.71 | 60.64 ±14.96 |
| – Theta | 61.42 ±11.67 | 65.44 ±15.03 |
| – Alpha | 61.18 ±11.95 | 65.41 ±14.86 |
| – Beta | 62.25 ±12.42 | 66.27 ±15.56 |

Removal of the **theta band** did not affect decoding performance. Accuracy remained stable at 71.53%, and AUC slightly increased to 79.30% (ΔAccuracy = −0.02 pp; ΔAUC = +0.012 relative to baseline). These differences were not statistically significant(Accuracy: t(19) = 0.029, p = 0.998; AUC: t(19) = −0.160, p = 0.998).

Similarly, removing the **alpha band** produced negligible changes in performance. Accuracy was 71.52% and AUC 79.18% (ΔAccuracy = −0.03 pp; ΔAUC ≈ 0.0000 relative to baseline), with no significant effects (Accuracy: t(19) = 0.102, p = 0.998; AUC: t(19) = −0.003, p = 0.998).

Finally, removing the **beta band** resulted in minimal changes. Accuracy decreased slightly to 71.30% and AUC to 79.03% (ΔAccuracy = −0.025 pp; ΔAUC = −0.015 relative to baseline), with non-significant paired comparisons (Accuracy: t(19) = 1.395, p = 0.477; AUC: t(19) = 0.649, p = 0.998).Overall, the inter-subject ablation analysis indicates that delta-band activity provides the primary contribution to cross-participant decoding, whereas removing higher-frequency bands (theta, alpha, beta) does not significantly impact classification performance under the LOSO framework.

**Table 8.**
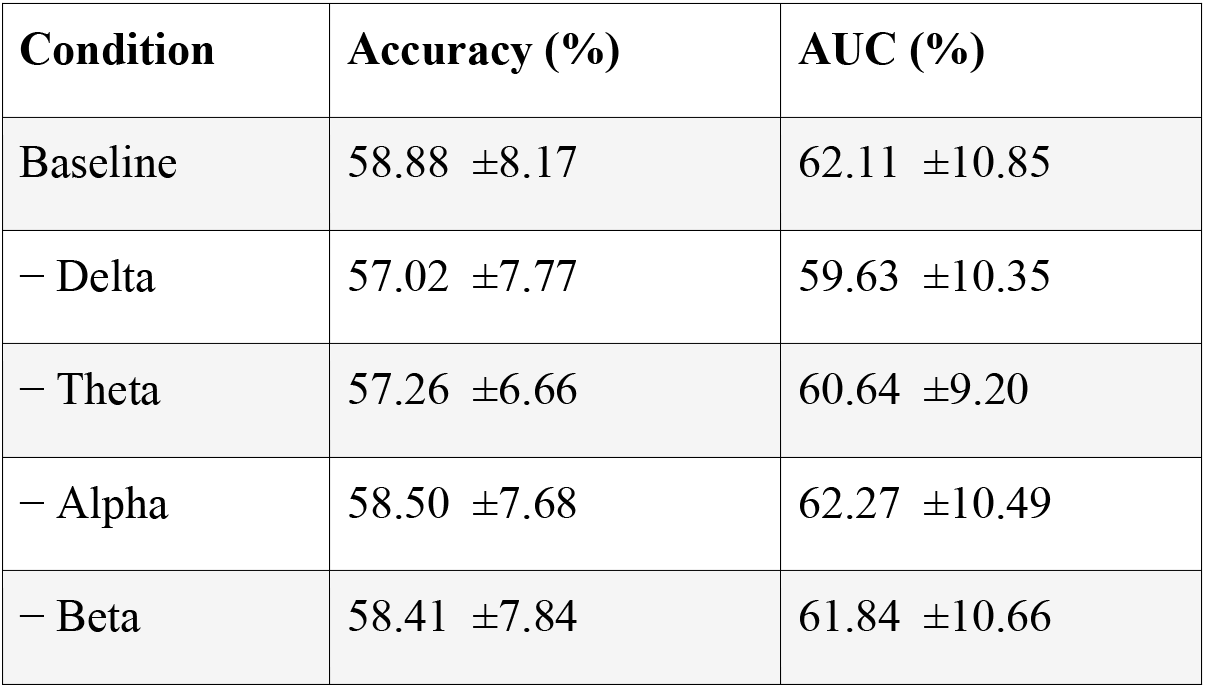
Inter-subject ablation analysis in Experiment 2 under a leave-one-subject-out (LOSO) evaluation scheme (N=17). Mean decoding Accuracy and AUC are reported for the baseline model and for band-ablation models in which one canonical frequency band (delta, theta, alpha, or beta) was removed at a time. Performance was computed on strictly held-out test participants and averaged across five runs using different validation participants to obtain participant-level estimates. Values are expressed as mean ±standard deviation across participants.

#### 5.4.2 Experiment 2: Oscillatory ablation impact on decoding System explainability

##### Intra-subject

Baseline performance (full spectrum) yielded a mean test accuracy of 62.5 ±12.0 % and a mean AUC of 66.1 ±15.2 % across participants.

**Delta-band** ablation produced the largest reduction in decoding performance. Removing the delta band decreased accuracy to **57.5** ±**11.7%** and AUC to **60.6** ±**15.0%**, corresponding to ΔAccuracy = **−5.0 pp** and ΔAUC = **−5.5 pp** relative to baseline. Paired t-tests confirmed that these reductions were statistically significant (Accuracy: t(16) ≈ **4.56**, p ≈ **0.0023**; AUC: t(16) ≈ **4.21**, p ≈ **0.0023**), with large effect sizes (Cohen’s d ≈ **1.11** for accuracy and ≈ **1.02** for AUC), indicating a dominant contribution of delta-band activity to decoding performance.

**Theta band** ablation produced only a modest decrease in performance. Accuracy reached **61.4** ±**11.7%**, and AUC **65.4** ±**15.0%**, corresponding to ΔAccuracy = **−1.1 pp** and ΔAUC = **−0.7 pp** relative to baseline. These differences did not remain significant after FDR correction (Accuracy: t(16) ≈ **2.04**, p ≈ **0.117**; AUC: t(16) ≈ **1.29**, p ≈ **0.285**), indicating a limited contribution of theta-band activity.

**Alpha band** ablation resulted in a small decrease in performance. Mean accuracy was **61.2** ±**12.0%**, and mean AUC **65.4** ±**14.9%**, corresponding to ΔAccuracy = **−1.3 pp** and ΔAUC = **−0.7 pp** relative to baseline. The reduction in accuracy reached statistical significance (Accuracy: t(16) ≈ **4.09**, p ≈ **0.0023**), whereas the difference in AUC did not(t(16) ≈ **1.80**, p ≈ **0.146**). This pattern suggests a limited and metric-dependent contribution of alpha-band activity.

Finally, **beta-band** ablation did not significantly affect decoding performance. Accuracy averaged **62.3** ±**12.4%**, and AUC **66.3** ±**15.6%**, corresponding to ΔAccuracy = **−0.3 pp** and ΔAUC = **+0.2 pp** relative to baseline. Paired t-tests confirmed the absence of a significant effect (Accuracy: t(16) ≈ **0.52**, p ≈ **0.697**; AUC: t(16) ≈ **−0.39**, p ≈ **0.702**).

Overall, these results indicate that **delta-band activity provides the dominant spectral contribution to intra-subject decoding performance in Experiment 2**. In contrast, **theta-and beta-band activity contribute minimally**, while **alpha-band effects remain weak and inconsistent across metrics**, suggesting that low-frequency dynamics constitute the primary source of discriminative information under the present decoding pipeline.

##### Inter-Subject

Baseline performance under the LOSO framework reached a mean accuracy of **58.9%** and an **AUC** of **62.1%** across participants.

Removing the **delta band** produced the largest degradation in decoding performance. Accuracy decreased to **57.0%** and AUC to **59.6%** (ΔAccuracy = **−1.9 pp**; ΔAUC = **−2.5 pp** relative to baseline). Paired t-tests confirmed that these reductions were statistically significant after FDR correction (Accuracy: t(16) = **2.85**, p = **0.0441**; AUC: t(16) = **2.68**, p = **0.0441**), indicating that low-frequency activity contributes substantially to neural representations that generalize across participants.

Removal of the **theta band** also led to a modest decrease in decoding performance. Accuracy reached **57.3%** and AUC **60.6%** (ΔAccuracy = **−1.6 pp**; ΔAUC = **−1.5 pp** relative to baseline). The reduction in accuracy remained significant after FDR correction (Accuracy: t(16) = **3.04**, p = **0.0441**), whereas the AUC difference did not survive correction (AUC: t(16) = **2.31**, p = **0.0686**), suggesting a limited and metric-dependent contribution of theta-band activity.

In contrast, removing the **alpha band** had a negligible impact on performance. Accuracy averaged **58.5%** and AUC **62.3%** (ΔAccuracy = **−0.4 pp**; ΔAUC = **+0.2 pp** relative to baseline), with no evidence for a reliable effect after correction (Accuracy: t(16) = **0.79**, p = **0.588**; AUC: t(16) = **−0.27**, p = **0.793**).

Similarly, removing the **beta band** did not significantly affect decoding performance. Accuracy averaged **58.4%** and AUC **61.8%** (ΔAccuracy = **−0.5 pp**; ΔAUC = **−0.3 pp** relative to baseline), with non-significant paired comparisons (Accuracy: t(16) = **0.99**, p = **0.543**; AUC: t(16) = **0.58**, p = **0.654**).

Overall, the inter-subject ablation analysis revealed a consistent spectral hierarchy with the intra-subject results. Delta-band activity provided the dominant contribution to decoding performance, while theta-band activity contributed modestly and inconsistently across metrics, and alpha and beta bands did not appear necessary for reliable cross-participant decoding under the LOSO framework.

### 5.5 Time-Frequency Dynamics of Agency-Related Neural Activity

To further characterize the neural dynamics underlying decoding performance, we conducted time-frequency analyses of EEG activity in the pre-auditory stimulus interval (−280 to +280 ms relative to keypress in Experiment 1 and from −560 to +280 ms in Experiment 2). This analysis aimed to identify frequency-specific neural signatures associated with variations in perceived agency.

Power estimates were extracted within canonical EEG frequency bands (delta: 2–4 Hz; theta: 4–8 Hz; alpha: 8–12 Hz; beta: 12–30 Hz) and compared between conditions using paired two-tailed t-tests at the participant level. To control for multiple comparisons across frequency bands, p-values were corrected using the Benjamini-Hochberg false discovery rate (FDR) procedure.

#### 5.5.1 Oscillatory Signatures of Agency under Decision Authority

Time–frequency analyses revealed significant differences between the Motor and AI without explanation conditions during the peri-action interval of interest. The strongest effects were observed in the low-frequency range.

**Figure 4.**
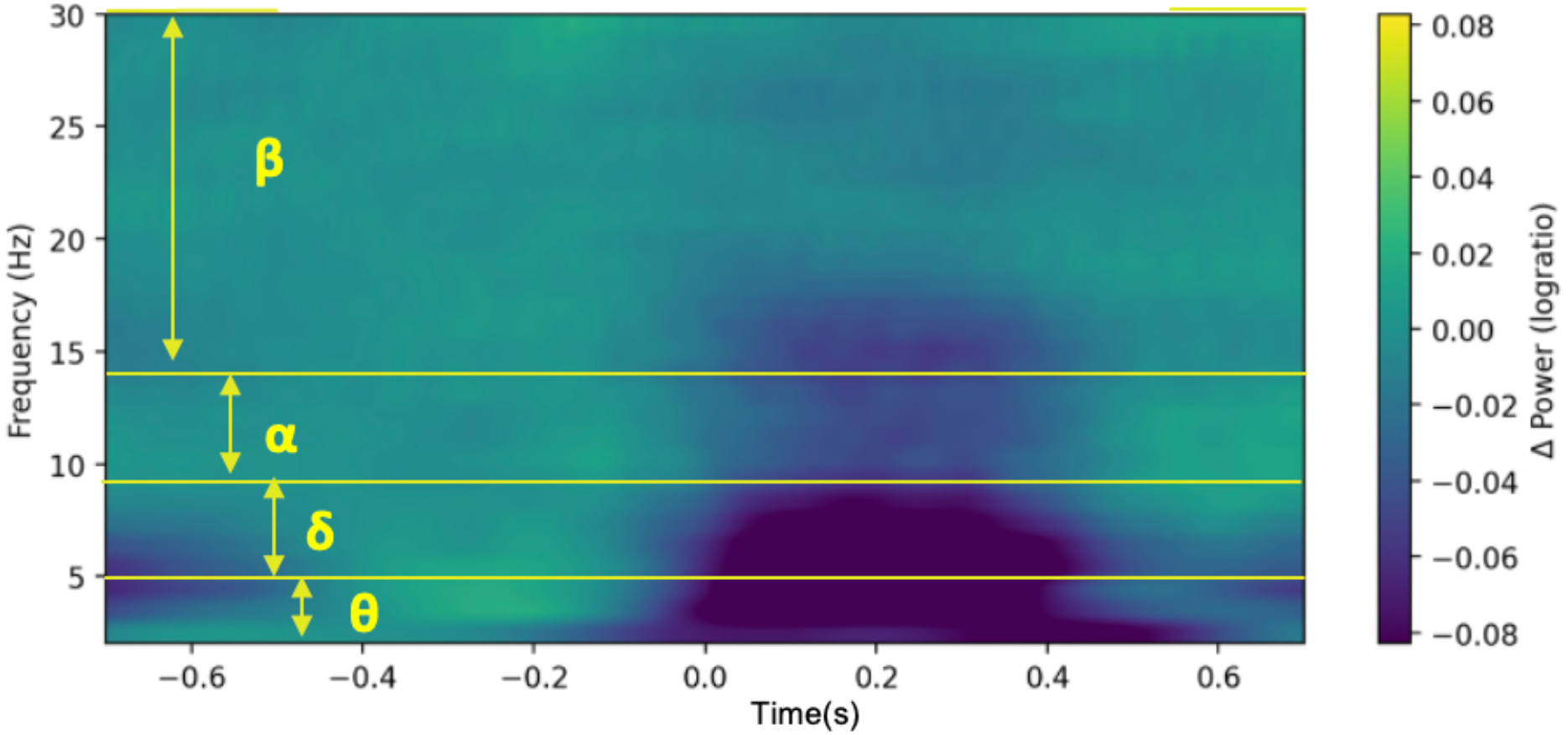
Time–frequency representation (TFR) of the difference in oscillatory power between the Motor and AI without explanation conditions in Experiment 1. TFRs were computed on extended epochs spanning −0.7 to +0.7 s relative to keypress for visualization purposes, allowing stable estimation of low-frequency activity and baseline normalization. Power is expressed as log-ratio change relative to baseline. The displayed map represents the participant-averaged condition contrast (Motor − AI without explanation). Statistical analyses were restricted to the pre-feedback interval used for EEGNet decoding, corresponding to the optimal temporal window identified by grid search (−0.280 to +0.280 s relative to keypress). Within this interval, significant differences were observed primarily in the delta and theta frequency bands. Visually, the strongest contrasts were concentrated in the post-keypress, pre-feedback portion of the analyzed window.

In the delta band, power was significantly lower in the Motor condition (M = −0.044) than in the AI without explanation condition (M = 0.024), t(19) = −3.101, p = 0.0118, Cohen’s d = −0.693. A comparable effect was observed in the theta band, with lower power in the Motor condition (M = −0.065) relative to the AI without explanation condition(M = −0.005), t(19) = −3.258, p = 0.0118, Cohen’s d = −0.729. No significant differences were observed in the higher-frequency bands after correction for multiple comparisons. In the alpha band, power was numerically lower in the Motor condition (M = −0.091) than in the AI without explanation condition (M = −0.067), but this effect did not reach significance, t(19) = −1.456, p = 0.2157, Cohen’s d = −0.326. Likewise, no reliable difference was observed in the beta band, t(19) = −1.191, p = 0.2483, Cohen’s d = −0.266.

Overall, these results indicate that differences associated with decision authority were accompanied by reliable modulations of delta- and theta-band activity during the peri-action interval, whereas no robust effects were observed in the alpha or beta ranges.

#### 5.5.2 Oscillatory Signatures of Agency under System Explainability

A similar pattern was observed when agency-related conditions were manipulated through system explainability. During the peri-action interval of interest (−0.560 to +0.280 s relative to keypress), significant differences emerged between the AI with explanation and AI without explanation conditions in the low-frequency range.

**Figure 5.**
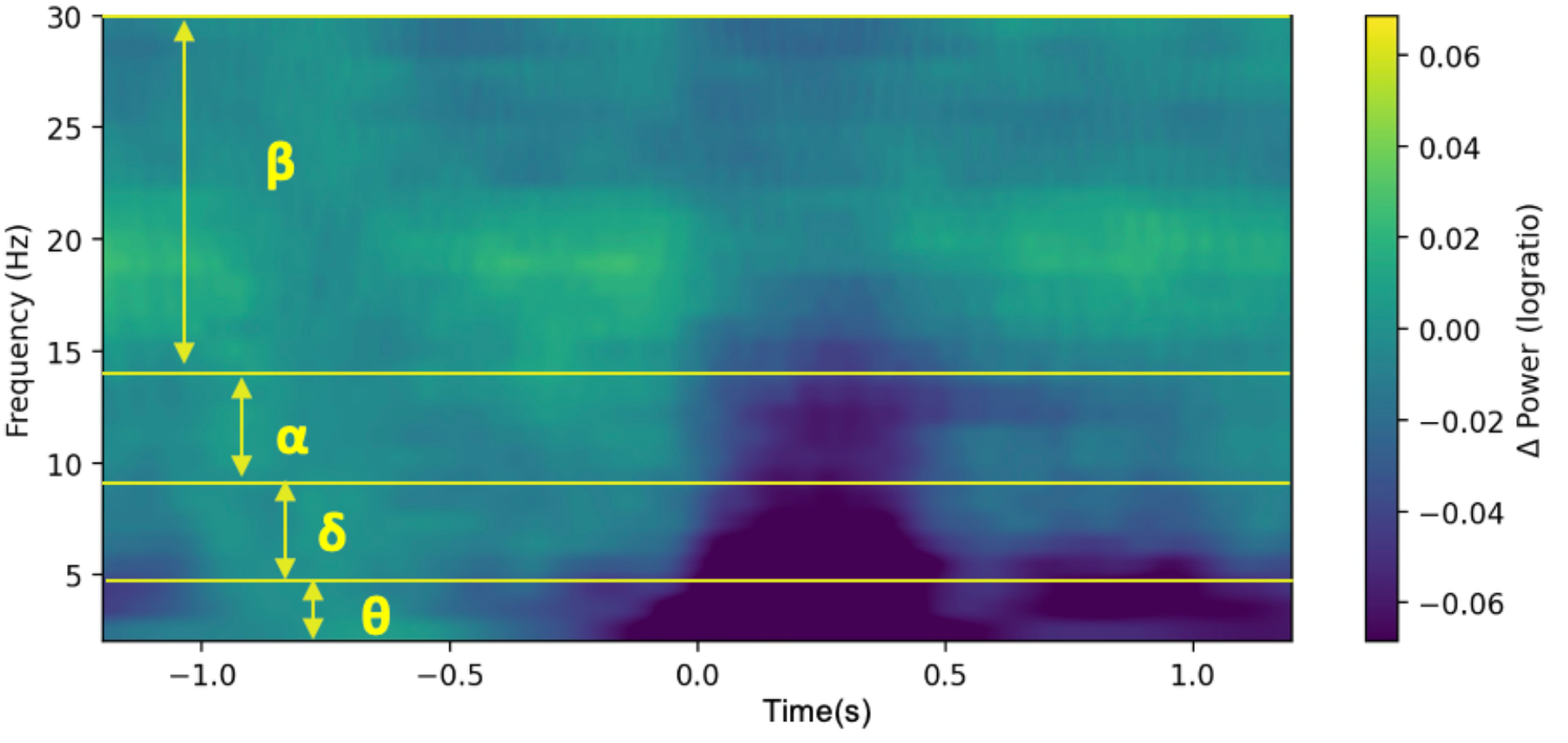
Time–frequency representation (TFR) of the difference in oscillatory power between the AI with explanation and the AI without explanation conditions in Experiment 2. TFRs were computed on extended epochs spanning −1.2 to +1.2 s relative to keypress for visualization purposes, allowing stable estimation of low-frequency activity and baseline normalization. Power is expressed as a log-ratio change relative to baseline. The displayed map represents the participant-averaged condition contrast (AI with explanation − AI without explanation). Statistical analyses were restricted to the pre-feedback interval used for EEGNet decoding, corresponding to the optimal temporal window identified by grid search (−0.560 to +0.280 s relative to keypress). Within this interval, significant differences were observed primarily in the delta and theta frequency bands. Visually, the strongest contrasts were concentrated in the post-keypress, pre-feedback portion of the analyzed window.

In the delta band, power was significantly lower in the AI with explanation condition (M = −0.043) than in the AI without explanation condition (M = 0.022), t(16) = −3.667, p = 0.0083, Cohen’s d = −0.889. A similar effect was observed in the theta band, with lower power in the AI with explanation condition (M = −0.043) relative to the AI without explanation condition (M = −0.002), t(16) = −2.805, p = 0.0254, Cohen’s d = −0.680.

By contrast, no significant differences survived correction in the higher-frequency bands. In the alpha band, power was numerically lower in the AI with explanation condition (M = −0.044) than in the AI without explanation condition (M = −0.025), but this effect did not reach significance after FDR correction, t(16) = −2.002, p = 0.0835, Cohen’s d = −0.485. Likewise, no reliable difference was observed in the beta band, t(16) = −0.971, p = 0.3461, Cohen’s d = −0.235.

Together, these results indicate that system explainability was associated with reliable modulations of delta- and theta-band activity during the peri-action interval, whereas no robust effects were observed in the alpha or beta ranges.

## 6. Discussion

The present study investigated whether agency-related states arising during human–AI interaction can be decoded from ongoing EEG activity and whether system explainability modulates their neural expression. Across two experiments, converging evidence was obtained at behavioral, decoding, spectral, and time-frequency levels. Subjective ratings confirmed that both decision authority and system explainability reliably modulated participants’ sense of control. At the neural level, deep-learning-based decoding demonstrated that these agency-related conditions could be discriminated from pre-feedback EEG activity both within individuals and, to a lesser extent, across individuals. Spectral ablation analyses further revealed a dominant contribution of low-frequency activity, particularly in the delta range, to decoding performance. Finally, time–frequency analyses showed that the most prominent neural differences were concentrated in the post-keypress, pre-feedback interval, especially within delta and theta bands. Taken together, these results demonstrate that agency-related neural states can be decoded from ongoing EEG activity before outcome feedback, indicating that agency-related information is expressed in predictive, pre-feedback neural dynamics rather than solely in post-outcome evaluation.

A central contribution of this work is the demonstration that agency-related states can be recovered from ongoing EEG activity without relying on discrete action–outcome events or trial-averaged event-related potentials. Traditional approaches to agency have largely depended on explicit self-reports or on event-locked neural measures such as intentional binding or ERP components, which are inherently limited in continuous interaction contexts (Wen et al., 2018). By contrast, the present results show that neural processes that covary with agency manipulations are embedded in single-trial EEG activity and can be recovered from ongoing signals. Importantly, achieving reliable decoding from single-trial EEG is particularly challenging due to its low signal-to-noise ratio, suggesting that the model captures structured and robust neural patterns rather than noise-driven fluctuations. These findings support the view that agency-related processes unfold dynamically over time, rather than emerging solely at discrete moments of action–outcome evaluation.

The **behavioral results** provide critical context for interpreting the neural findings. In Experiment 1, delegating decision authority to the AI significantly reduced perceived control, consistent with prior work showing that increased automation attenuates the sense of agency (Endsley & Kiris, 1995; Kaber & Endsley, 1997; Berberian, 2019; Houdoyer et al., under review). In Experiment 2, providing distal explanations significantly increased perceived control, in line with studies demonstrating that intention-based explainability can partially restore agency in human–AI interaction (Le Goff et al., 2018; Vantrepotte et al., 2022; Houdoyer et al., under review). These effects confirm that the experimental manipulations induced meaningful variations in subjective control, supporting the interpretation that decoding performance reflects differences in agency-related processing rather than arbitrary task conditions.

At the neural level, **EEGNet decoding performance** indicates that these agency-related states are reliably encoded in EEG activity. Intra-subject decoding yielded robust performance (Experiment 1: Accuracy: 77.61 ±9.17% / AUC: 84.25 ± 9.73%; Experiment 2: Accuracy: 62.50 ±11.98% / AUC: 66.11 ± 15.19%) in both experiments (for details see Tables 1 & 3 and Figures 2 & 3), demonstrating stable condition-related signal signatures at the individual level. Cross-subject decoding remained significantly above chance (Experiment 1: Accuracy: 71.55 ± 8.09% / AUC: 79.18 ± 8.23%; Experiment 2: Accuracy: 58.88 ± 8.17% / AUC: 62.11 ± 10.85%; see Tables 2 & 4), indicating the presence of partially shared neural structure across participants despite inter-individual variability. Such generalization is particularly important in the context of human-centered AI, where scalable systems require neural markers that extend beyond subject-specific calibration. At the same time, decoding performance differed between the two experimental manipulations. Classification was consistently stronger when agency was manipulated through decision authority than when it was modulated through system explainability. This dissociation suggests that direct control over action selection gives rise to more robust and consistently expressed neural signatures than interpretability-based modulation. This distinction is theoretically informative, as it indicates that different components of agency, control-based versus inference-based, may rely on partially distinct neural mechanisms or may differ in the strength and consistency of their neural expression (see also Weiss et al., 2022). While both manipulations influence subjective experience, their neural correlates may differ in their accessibility to EEG-based decoding.

**Spectral ablation analyses** provide further insight into the structure of these neural representations. When focusing on decoding accuracy, a clear hierarchy emerged across both experiments, with delta-band activity consistently providing the strongest contribution to decoding performance. Removal of the delta band systematically produced the largest and most reliable decrease in accuracy, both in intra-subject and inter-subject decoding. This convergence across experiments and evaluation regimes indicates that slow neural dynamics carry a substantial proportion of the discriminative information exploited by the model.

Beyond this dominant delta contribution, the pattern observed for theta-band ablation differed between the two experiments. In Experiment 1, removing theta activity significantly reduced intra-subject decoding accuracy, suggesting that theta dynamics, in addition to delta activity, contributed to the participant-specific representation of decision authority. By contrast, in Experiment 2, theta-band ablation did not significantly affect intra-subject decoding, but did reduce inter-subject decoding accuracy, indicating a weaker yet more generalizable contribution of theta activity when agency was modulated through system explainability. This dissociation suggests that theta-band activity may play different roles depending on the nature of the agency manipulation: under direct control, it may contribute primarily to subject-specific representations, whereas under explanation-based modulation, it may reflect weaker but more shared processes across participants.

This difference likely reflects the distinct cognitive demands associated with the two experimental manipulations. In Experiment 1, agency was manipulated through direct control over action selection, a condition that strongly engages performance monitoring and cognitive control processes. Theta-band activity, particularly over fronto-midline regions, has been consistently associated with such functions, including action monitoring and adaptive control. In this context, theta dynamics may capture participant-specific monitoring strategies, explaining their contribution to intra-subject decoding but limited generalization across individuals. By contrast, in Experiment 2, agency was modulated through system explainability, a manipulation that relies less on direct action control and more on the formation of internal models of the system’s behavior. Such processes are likely more homogeneous across participants and less dependent on individual control strategies, resulting in weaker intra-subject effects but modest contributions to cross-participant generalization.

Alpha-band ablation showed a limited and context-dependent effect, reaching significance only for intra-subject decoding in Experiment 2, whereas beta-band ablation had no measurable impact on decoding accuracy across analyses.

Overall, this pattern indicates that slow neural dynamics, and particularly delta-band activity, carry the most informative signal for decoding agency-related states, while theta activity plays a secondary and manipulation-dependent role. This interpretation is consistent with previous work linking low-frequency oscillations to large-scale integration and predictive processing (Cavanagh & Frank, 2014; Arnal & Giraud, 2012). Importantly, the hierarchy observed here, with delta as the dominant contributor and other bands showing weaker or more context-dependent effects, suggests that agency-related information is preferentially encoded at the slowest temporal scales in the present task in the present paradigm.

Notably, no robust contribution of alpha–mu (Kang et al., 2015; Wen et al., 2017) or beta-band activity was observed, despite their frequent association with sensorimotor aspects of agency.

This absence does not contradict prior findings, but rather suggests that under the present task constraints, characterized by limited motor demands and a focus on anticipatory processes, the most informative neural signals were carried by slower dynamics related to monitoring and prediction.

**Time–frequency analyses** further clarify both the temporal and functional characteristics of these effects. The most salient differences between conditions were visually concentrated in the post-keypress, pre-feedback interval, indicating that the decoded signal primarily reflects anticipatory and predictive processes occurring after action initiation and before outcome perception. This temporal profile is consistent comparator models of agency, in which internal predictions about action outcomes are generated and evaluated prior to feedback (Frith et al., 2000; Blakemore et al., 2002; Haggard, 2017). The present findings extend these models by demonstrating that such predictive processes can be detected from ongoing EEG activity and contribute directly to decoding performance. Crucially, time–frequency analyses revealed that delta- and theta-band power were significantly higher in the AI without explanation condition than in both the Motor condition (Experiment 1) and the AI with explanation condition (Experiment 2). Thus, conditions associated with reduced perceived agency consistently exhibited increased low-frequency power in the pre-feedback interval. This pattern provides a mechanistic link between decoding, ablation, and time–frequency results, suggesting that the classifier relied on systematic increases in low-frequency activity associated with heightened monitoring demands or reduced predictability.

From a neurophysiological perspective, increased delta and theta activity has been associated with performance monitoring, cognitive control, and the processing of uncertainty or mismatch between expected and actual outcomes (Cavanagh & Frank, 2014; Arnal & Giraud, 2012). In the present context, higher low-frequency power in the AI-without-explanation condition may reflect increased monitoring demands or reduced predictability of system behavior when users lack access to the agent’s intentions. Conversely, reduced delta and theta power in the Motor and AI with explanation conditions may indicate more efficient predictive processing and reduced monitoring load, consistent with a stronger sense of control. Importantly, this pattern was consistent across both experimental manipulations. Conditions associated with higher perceived agency, namely direct motor control and AI-with-explanation, were both characterized by reduced low-frequency power relative to the AI without explanation condition. This convergence suggests that delta- and theta-band activity may index shared neural mechanisms subserving the monitoring of the unfolding interaction and the intermediate effects arising during its course, prior to the final outcome, under conditions of reduced control or increased uncertainty.

An important strength of the present study lies in the convergence between deep learning and classical neurophysiological analyses. Decoding, ablation, and time–frequency approaches independently identified low-frequency, pre-feedback neural dynamics as the primary source of discriminative information, strengthening the robustness of the findings. These results have important implications for neuroengineering applications. By demonstrating that agency-related information can be extracted from ongoing EEG activity, this work opens the possibility of real-time, non-intrusive monitoring of user states. Such capability could support adaptive human-AI systems that dynamically regulate automation or explanatory support to maintain an appropriate sense of control. The presence of above-chance inter-subject decoding further suggests that such approaches may generalize across users.

Several limitations should nevertheless be acknowledged. First, although the experimental manipulations were behaviorally validated, the present study does not establish a direct relationship between decoding performance and trial-wise subjective reports of agency. Due to the limited proportion of trials including behavioural ratings, such correlations could not be assessed. As a result, decoded signals should be interpreted not as direct readouts of subjective agency, but as reflecting **agency-related** computations embedded within broader cognitive functions, including attention, monitoring, predictive inference, and metacognitive evaluation. Such an interpretation aligns with multifactorial accounts of agency, according to which the sense of agency emerges from the integration of multiple sensorimotor, contextual, and cognitive cues rather than from any single process-specific mechanism (Synofzik et al., 2008; Chambon et al., 2014).

Second, inter-subject decoding performance, although significant, remained lower than intra-subject performance, highlighting the impact of inter-individual variability on the generalization of neural signatures of agency.

Third, alpha and beta frequency bands did not show reliable contributions in either the decoding analyses, as assessed by the EEGNet ablation procedure, or the time–frequency analyses. This finding contrasts with previous studies linking the sense of agency to sensorimotor alpha–mu and beta oscillations (Kang et al., 2015; Wen et al., 2017; Bu-Omer et al., 2021). One possible explanation is that the present paradigm primarily engaged cognitive and decisional aspects of agency, such as anticipation, delegation of control, and the interpretation of system behaviour, rather than continuous motor control. In such a context, agency-related processes may rely less on sensorimotor integration and more on predictive and inferential mechanisms. In addition, the decoding analysis focused on pre-feedback activity, thereby targeting anticipatory processes. Given that alpha–mu suppression is typically observed during or following movement execution and is closely linked to motor preparation and feedback monitoring (Wen et al., 2017), its contribution may have been reduced within the temporal window examined here. The relatively limited motor demands of the task may have further attenuated the involvement of canonical sensorimotor rhythms.

Finally, while the interpretation of post-keypress activity as reflecting outcome anticipation is theoretically grounded, it remains indirect and would benefit from more targeted experimental manipulations. Taken together, these limitations suggest that agency-related neural signatures may not rely on a single canonical marker, but rather emerge from the interaction between predictive, decisional, and control-related processes, whose relative contribution depends on task demands and temporal dynamics.

In conclusion, the present study demonstrates that agency-related neural states can be decoded from ongoing EEG activity before the occurrence of outcome feedback. This finding indicates that agency is not only reconstructed retrospectively from the sensory consequences of action, but also relies on earlier anticipatory and monitoring processes unfolding during action execution. The fact that decoding performance was significant not only within participants, but also across participants, further suggests that some EEG signatures of agency are shared across individuals rather than being purely idiosyncratic. This has important implications for the development of neuroadaptive systems, as it indicates that agency-related states may be tracked across users without requiring complete recalibration for each individual. The complementary oscillatory analyses provide further insight into the neural dynamics underlying this decoding. Agency-related information was primarily expressed in low-frequency activity, particularly in the delta and theta ranges, and was most prominent in the post-keypress, pre-feedback interval. Increased low-frequency activity was associated with reduced sense of agency, which may reflect greater uncertainty, enhanced monitoring demands, or less precise predictions about the consequences of action. Although these oscillatory findings remain exploratory, they support a dynamic account of agency in which perceived control emerges from ongoing predictive and monitoring processes rather than from post hoc outcome evaluation alone.

By integrating behavioural validation, deep learning-based EEG decoding, and oscillatory analyses, this work provides a unified framework for studying agency as a dynamic neural process. More broadly, it lays the groundwork for neuroadaptive systems capable of tracking fluctuations in human agency during interaction and, ultimately, of supporting users in contexts where control is partially delegated to automated or AI-based systems.

